# Efficacy of the β-lactam\β-lactamase inhibitor combination is linked to WhiB4 mediated changes in redox physiology of *Mycobacterium tuberculosis*

**DOI:** 10.1101/103028

**Authors:** Saurabh Mishra, Prashant Shukla, Ashima Bhaskar, Kushi Anand, Priyanka Baloni, Rajeev Kumar Jha, Abhilash Mohan, Raju S. Rajmani, V. Nagaraja, Nagasuma Chandra, Amit Singh

## Abstract

**Aims:** Inhibition of β-lactamase by clavulanate (Clav) sensitizes multi-and extensively drug-resistant *Mycobacterium tuberculosis* (*Mtb*) strains towards β-lactams such as amoxicillin (Amox). However, the underlying mechanism of how *Mtb* responds to Amox-Clav combination (Augmentin; AG) is not characterized.

**Results:** We integrated global expression profiling with the protein-protein interaction landscape and generated a genome-scale network of *Mtb* in response to AG. In addition to specific targets (*e.g.,* peptidoglycan biosynthesis and β-lactamase), the response to AG was also centered on redox-balance, central carbon metabolism (CCM), and respiration in *Mtb.* We discovered that AG modulates superoxide levels, NADH/NAD^+^ balance and mycothiol redox potential (*E_MSH_*) of *Mtb*. Higher intra-mycobacterial *E_MSH_* potentiates mycobactericidal efficacy of AG, whereas lower *E_MSH_* induces tolerance. Further, *Mtb* responds to AG via a redox-sensitive transcription factor, WhiB4. *MtbΔwhiB4* displayed higher expression of genes involved in β-lactam resistance along with those mediating respiration, CCM and redox balance. Moreimportantly, WhiB4 binds to the promoter regions and represses transcription of genes involved in β-lactamase expression in a redox-dependent manner. Lastly, while *MtbΔwhiB4* maintained internal *E_MSH_*, exhibited greater β-lactamase activity and displayed AG-tolerance, overexpression of WhiB4 induced oxidative shift in *E_MSH_* and repressed β-lactamase activity to aggravate AG-mediated killing of drug-sensitive and –resistant strains of *Mtb*.

**Innovation and Conclusions:** This work demonstrate that efficacy of β-lactam\β-lactamase inhibitor combination can be attenuated by elevating mycobacterial antioxidant capabilities and potentiated by impairing redox buffering capacity of *Mtb*. The functional linkage between β-lactams, redox balance, and WhiB4 can be exploited to potentiate AG action against drug-resistant *Mtb*.

## Introduction

*Mycobacterium tuberculosis* (*Mtb)* displays tolerance to several clinically important antibacterials such as aminoglycosides and β-lactams (14,36). Innate resistance of *Mtb* towards β-lactams is largely considered due to the presence of a broad-spectrum Ambler class A β- lactamase (BlaC) (15). Other mechanisms such as cell envelope permeability, induction of drug efflux pumps, and variations in peptidoglycan biosynthetic enzymes also play a role in β-lactam-resistance in *Mtb* (18,30). The Ambler class A β -lactamases are mostly susceptible to inhibition by clavulanate (Clav), sulbactam (Sub), and tazobactam (Taz) (30). Indeed, intrinsic resistance of *Mtb* towards β-lactams can be overcome by using the combination of β-lactams with Clav (5,22). Combination of amoxicillin (Amox) with Clav (Augmentin; AG) was active against *Mtb in vitro* and had significant early bactericidal activity in patients with drug-resistant TB (5,8). Likewise, combination of meropenem with Clav showed significant bactericidal activity against drug-resistant strains of *Mtb* (22). With this attention, there is an urgent need to generate knowledge about the mechanisms of action of β-lactams in combination with Clav against *Mtb* as well as possible route(s) of resistance.

In other bacteria, β-lactams directly interact with enzymes involved in peptidoglycan (PG) synthesis, which may lead to killing via multiple mechanisms including induction of autolysin pathway, holin:antiholin pathways, DNA damage, and alterations in physiology (*e.g.,* TCA cycle, and oxidative stress) (26,28,34,42,54). The complex effects of β-lactams on both PG biosynthesis and other processes indicate that the response to β-lactams could be mediated either through direct sensing of β-lactam molecules or by their effects on bacterial physiology. In *Staphylococcus aureus (S. aureus),* a transmembrane protease, BlaR1, senses β-lactam concentrations by direct binding through extracellular domain, which activates its intra-cytoplasmic proteolytic domain resulting in cleavage of the β-lactamase repressor, BlaI, and induction of β-lactamase expression (17). The knowledge of how β-lactams are sensed to activate adaptation programs is lacking in *Mtb*. It has been shown that *Mtb* expresses a close homolog of BlaR1 encoded by Rv1845c (*blaR*), which modulates the activity of BlaC by regulating BlaI repressor in a manner analogous to *S. aureus* BlaR1-BlaI couple (45). However, BlaR1 orthologues in all mycobacterial species lack extracellular sensor domain involved in binding with β-lactams (45), indicating that mechanisms of antibiotic sensing and BlaC regulation are likely to be distinct in *Mtb*. Similarly, how β-lactams influence mycobacterial physiology such as redox balance and metabolism remain unknown. Therefore, novel insights are needed to discover how the presence of β-lactams is conveyed in *Mtb* to activate appropriate adaptation response.

Here, we generated a system-scale understanding of how AG affects mycobacterial physiology. Exploiting a range of technologies, we explained mechanistically that AG efficacy is partly dependent upon the redox physiology of *Mtb*. Further, we described rationally the role of a redox-responsive transcription factor, WhiB4, in regulating tolerance to AG by recalibrating drug transcriptome, β-lactamase activity, and intramycobacterial mycothiol redox potential (*E_MSH_*) during infection. Our study demonstrates how *Mtb* alters its redox physiology in response to AG and identifies MSH and WhiB4 as major contributors to β-lactam tolerance.

## Results

### Network analysis revealed modulation of cell wall processes in response to AG in *Mtb*

To assess the response of *Mtb* towards β-lactam and β-lactamase inhibitor combination(s), we generated transcriptome of mycobacterial cells upon exposure to AG. We observed that 100 µg/ml of Amox in combination with 8 µg/ml of Clav (10X MIC of AG) arrested bacterial growth at 6 h and displayed killing only after 12 h post-exposure (Supplementary Fig. 1-Inset). Therefore, expression changes at a pre-lethal phase (*i.e.* 6 h post-10X MIC AG treatment) can reveal significant insights into pathways *Mtb* exploits to tolerate AG.

A total of 481 genes were induced (≥2-fold; *P* value ≤ 0.05) and 461 were repressed (≥2-fold; *P* value ≤ 0.05) in the wt *Mtb* upon AG-treatment (Supplementary Table S1). While important, transcriptome alone represent only a narrow glimpse of the mechanisms exploited by a cell in order to tolerate drug exposure. Therefore, it is important to harness the power of computational approaches that combine condition-specific expression data with general protein interaction data to construct dynamic and stress response networks. We first created a comprehensive protein-protein interactome (PPI) of *Mtb* using information from experimentally validated and published interactions (see *Materials and Methods*) and then integrated microarray data with the PPI to generate AG response network. In the network, a node represents a gene and node weight is the product of normalized intensity values with the corresponding fold change in expression upon drug exposure. Expression profiles of nodes were used to determine the edge-weight between two interacting nodes. Although a full description of the mathematical equations and algorithms used to generate AG response network is beyond the scope of this study, we encourage readers to refer our original papers for detailed methodology (39,46,47).

Supplementary Fig. 1 and Table S2 represent the 1 % top nodes, which covers a total of 806 genes which are connected through 1096 interactions to form a well-connected AG response network of *Mtb*. Genes belonging to diverse functional classes such as intermediary metabolism, cell wall, lipid metabolism, virulence, and information pathways featured in the response network. Since β–lactams primarily target PG-biosynthetic processes, our expression data clearly showed induction of genes associated with the biogenesis of PG and other cell processes (Fig. 1A). Network analysis confirmed that the cumulative node weight intensity (CNW) of genes belonging to cell wall related processes was highest (30616361.02) among the classes affected by AG treatment (Fig. 1B). Induction of genes involved in PG biosynthesis and β-lactamase regulation (*blaR-blaI*) (Fig. 1A) in response to AG validates our experimental conditions. In addition, two other mechanisms involved in tolerating β-lactams *i.e.* outer membrane permeability (mycolic acid biogenesis [*kasA, kasB,* and *fabD*] and *omp*) and efflux pumps (*efpA, Rv1819c* and *uppP*) were induced in response to AG (Supplementary Table S1). Importantly, in line with cell surface targeting activity of β-lactams, our analysis identified activation of global regulators (*e.g., sigE*, *sigB, and mprAB*) of cell envelope stress in *Mtb*. These regulators function as major hub nodes and form-interconnected networks of genes important to maintain cell wall integrity in response to AG (Fig. 2). Altogether, *Mtb* responds to AG by modulating the expression of cell envelope associated pathways including those that are the specific targets of β-lactams.

**Fig 1:**
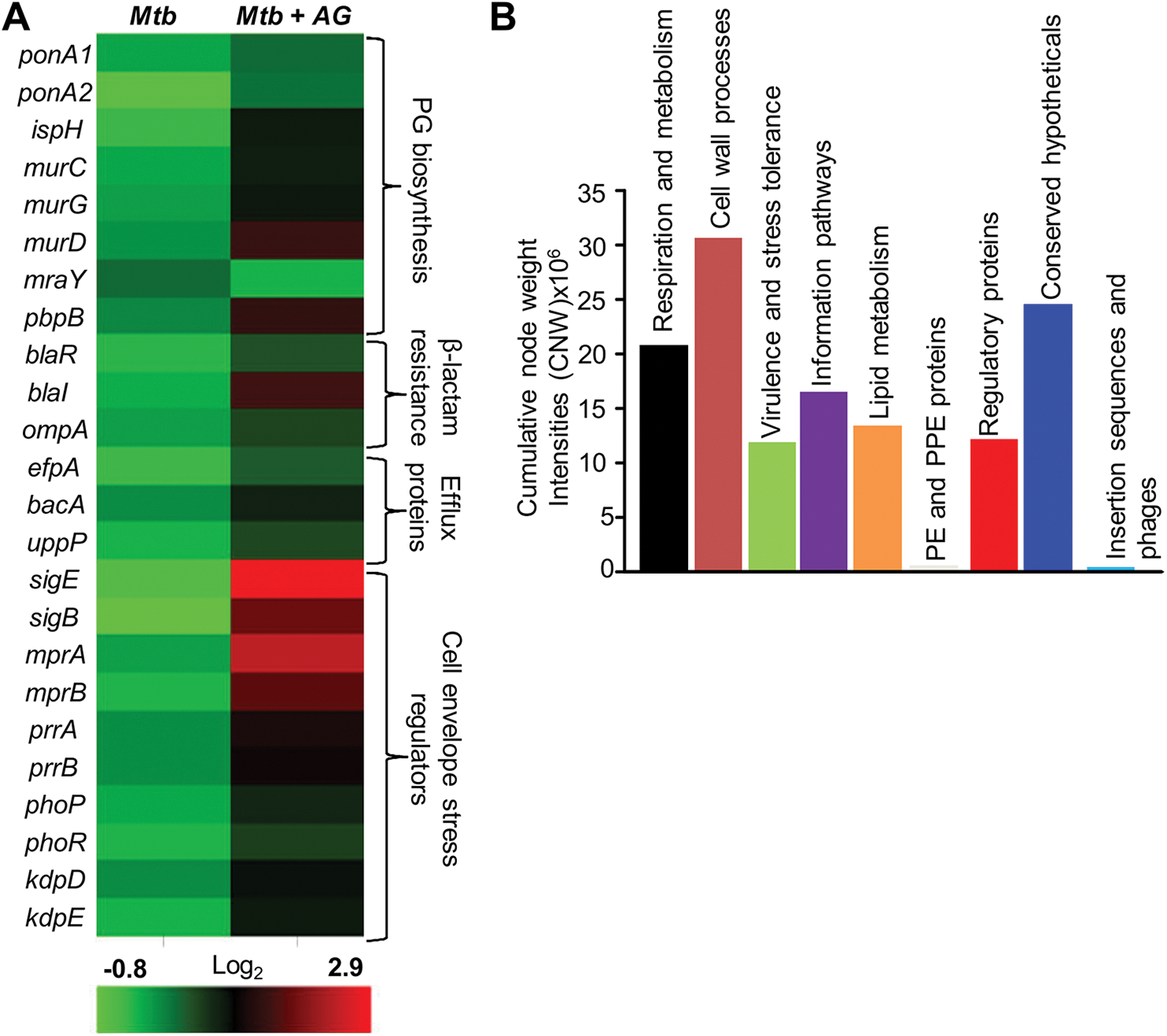
Network analysis identified pathways affected by AG exposure in *Mtb*. Wt *Mtb* was grown to an OD_600_ of 0.4 and treated with 100 µg/ml of Amox and 8 µg/ml of Clav (AG) for 6 h at 37°C. Total RNA was isolated and processed for microarray analysis as described in *Materials and Methods*. (A) Heat map shows the fold-change (log_2_) in expression of genes belonging to cell wall processes for untreated and 6 h of AG-treatment of *Mtb* from two biological samples. (B) Cumulative node weight intensities (CNW) were derived by addition of the node weights of genes in a particular functional group upon exposure to AG. Node weight intensity of a gene was derived by multiplying the normalized intensity value with the corresponding fold-change (FC) value.

**Fig 2:**
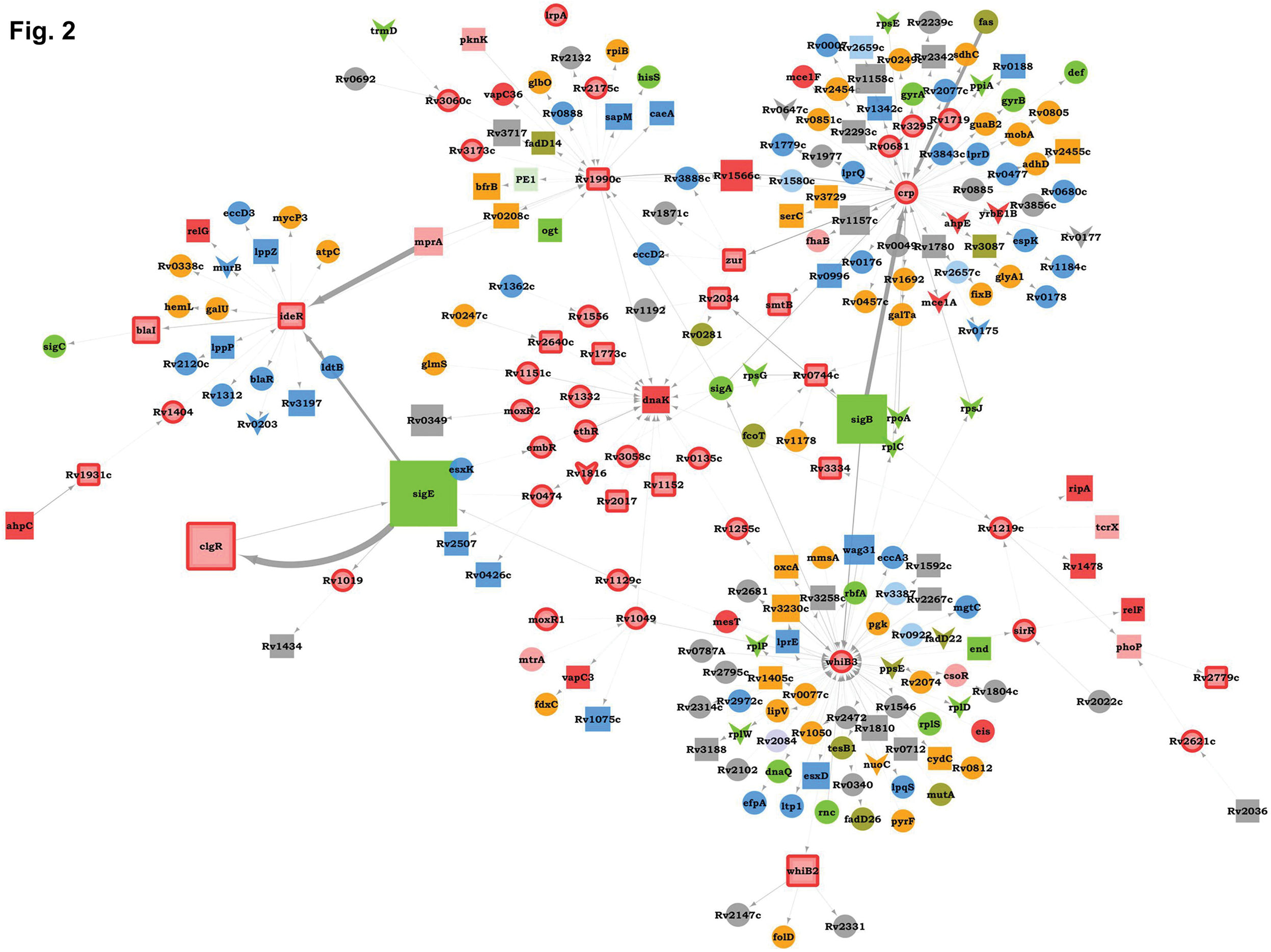
Sub-network of major hub nodes showing the top-most activities regulating response of *Mtb* upon AG treatment. The nodes are colored according to the functional modules they belong to, and edge thickness reflects the strength of the interaction. Shapes of the nodes denotes regulation of the gene expression (square: induced; arrowhead: repressed; and circle: constitutive). Functional modules based on annotations given in the TubercuLlist (http://tuberculist.epfl.ch/) include red: virulence, detoxification, and adaptation, blue: cell wall and cell processes, green: information pathways, orange: intermediary metabolism and respiration, olive green: lipid metabolism, grey: conserved hypotheticals, pink: regulatory proteins, cyan: insertion sequences and phages, and light green: PE/PPE family.

### AG affects pathways associated with CCM, respiration, and redox balance

Our analysis revealed enrichment of nodes belonging to “intermediary metabolism and respiration” within AG response network (CNW= 20716788.92; Fig. 1B). For example, energetically efficient respiratory complexes such as NADH dehydrogenase I (*nuo* operon) and ATP-synthase (*atpC, atpG, atpH*) were down-regulated, whereas energetically less favored NADH dehydrogenase type II (*ndh*), cytochrome bd oxidase (*cydAB*), and nitrite reductase (*nirBD*) were activated in response to AG (Fig. 3A). AG also substantially induced genes involved in assembly and maturation of cytochrome bd oxidase in *E. coli* (*i.e. cydDC*; ABC transporter) (7) (Fig. 3A). The transcriptional shift towards a lesser energy state is consistent with the down-regulation of several genes associated with TCA cycle (*sucCD, fum, mdh, and citA*), along with an induction of glycolytic (*pfkA, pfkb, fba, and pgi*), gluconeogenesis (*pckA*) and glyoxylate (*icl1*) pathways (Fig. 3A). Interestingly, *icl1* has recently been shown to promote tolerance of *Mtb* towards diverse anti-TB drugs by maintaining redox homeostasis (37). These findings indicate that rather than energy generation, maintenance of redox balance is more likely to be an important cellular strategy against AG.

**Fig 3:**
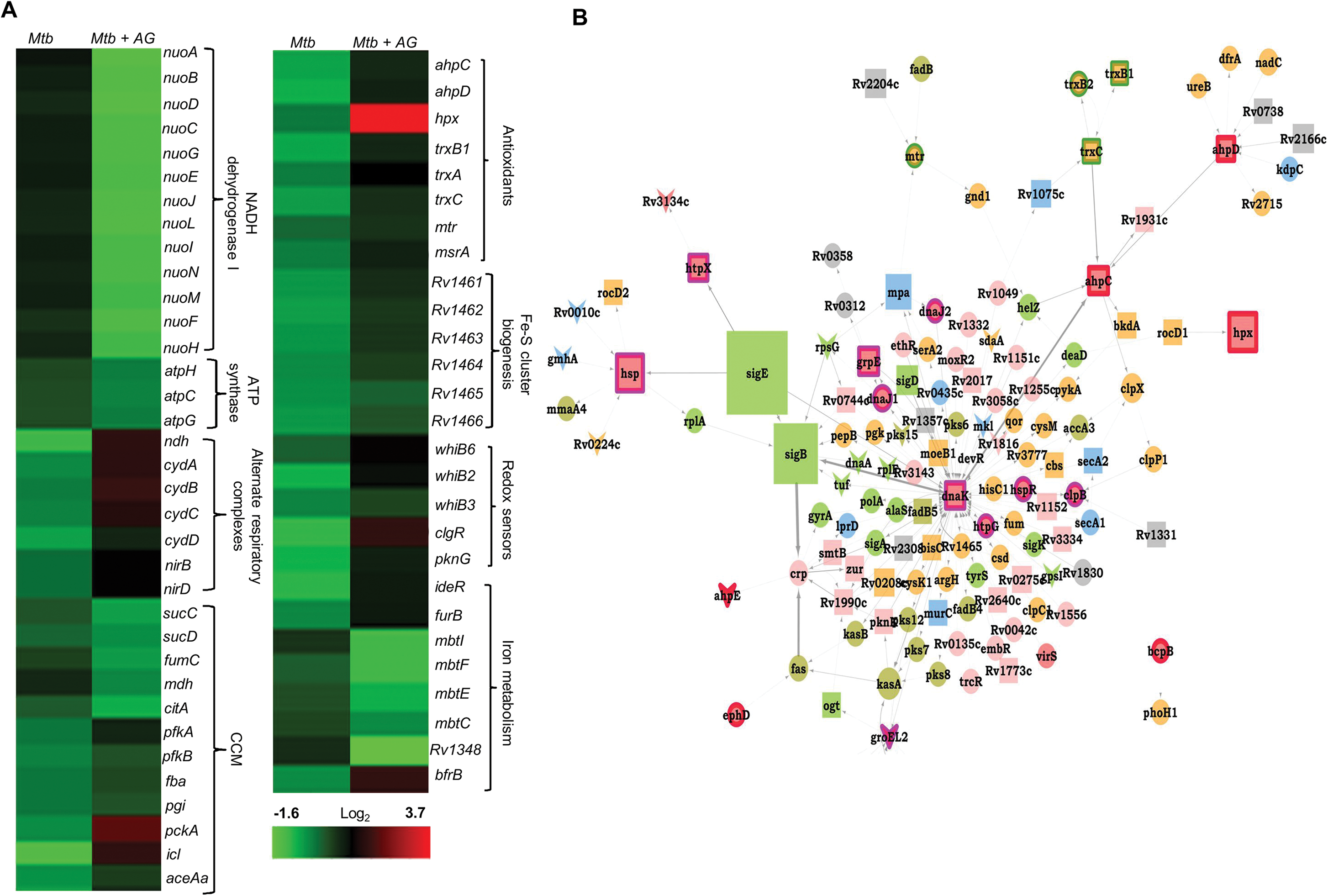
AG influences multiple pathways involved in central metabolism, respiration and redox balance in *Mtb*. (A) Heat maps depicting fold-change (log_2_) in expression of genes coordinating CCM, respiration, and redox balance for untreated and 6 h of AG-treated *Mtb* from two biological samples. (B) Sub-network showing close interactions between diverse regulators of oxidative stress including stress-responsive sigma factors, antioxidants, chaperones, and redox-sensitive transcription factors during AG treatment. Description of nodes function, color codes, edge thickness, and shape of nodes is as given in figure 1.

The influence of AG on mycobacterial redox physiology is also apparent by a significant enrichment of genes that can report exposure to oxidative stress in the response network. We found increased expression of reactive oxygen species (ROS) detoxifying enzymes (*ahpCD, katG,* and *hpx*), antioxidant buffers (*trxB1, trxA, trxC, mtr*), methionine sulfoxide reductase (*msrA*), Fe-S cluster repair system (Rv1461-Rv1466; *suf*), and intracellular redox sensors (*whiB6*, *whiB2, whiB3, and pknG)* (Fig. 3A). The global regulator of oxidative stress in bacteria, OxyR, is non-functional in *Mtb* (9). However, we earlier reported that a redox-sensitive DNA binding protein, WhiB4, functions as a negative regulator of OxyR-specific antioxidant genes (*e.g., ahpCD*) in *Mtb* (6). Consequently, *Mtb* lacking *whiB4* (*MtbΔwhiB4*) displayed higher expression of antioxidants and greater resistance towards oxidative stress (6). While microarrays in response to AG showed only a modest repression of WhiB4 (~1.3 fold), our qRT-PCR under similar conditions confirmed a significant down-regulation (-5.00 ± 0.27 fold; *P* value ≤ 0.001) as compared to unstressed *Mtb*. The breakdown of iron homeostasis is another hallmark of oxidative stress (23). Data showed induction of Fe-responsive repressors, *ideR* and *furB*, along with the down-regulation of genes encoding Fe-siderophore biosynthetic enzymes (*mbt* operon) and Fe-transport (*Rv1348*), and up-regulation of Fe-storage (*bfrB*) (Fig. 2A). Studies have suggested an important role for DnaK and ClpB chaperones in promoting recovery from oxidative stress (12,57). Our analysis identified that most of the functionally diverse nodes (sigma factors, antioxidants and redox-sensors) converge at a common stress responsive chaperone, DnaK, making it a major hub node coordinating AG stress response in *Mtb* (Fig. 3B).

Recently, two mycobacterial redox buffers, mycothiol (MSH) and ergothioniene (EGT), were implicated in protecting against oxidants and antibiotics (44). We compared gene expression changes displayed by MSH and EGT mutants (44) with the AG transcriptome. An ~ 60% of genes regulated by MSH and EGT also displayed altered expression in response to AG (Supplementary Fig. S2), indicating an overlapping roles of MSH and EGT in tolerating oxidative stress and AG in *Mtb* (44). Lastly, we performed transcriptomics of *Mtb* in response to a known oxidant cumene hydroperoxide (CHP; 250 μM for 1 h [non-toxic concentration]) and compared expression changes with AG-responsive regulons. As shown in Supplementary Fig. S3, a considerable overlap in gene expression (~30%) was observed between the two conditions (Supplementary Table S4). More importantly, genes associated with β-lactams tolerance (*ponA2*, *ispH,* and *kasA*) and redox-metabolism (*ahpCD*, *trxB1*, *trxB2*, *trxC*, and *suf*) were similarly regulated under CHP and AG challenge (Supplementary Table S4). Lastly, we validated our microarray data by performing qRT-PCR on a few genes highly deregulated upon AG treatment (Supplementary Table S3). Taken together, data indicate a major recalibration of genes regulating mycobacterial redox physiology in response to AG.

### AG treatment induces redox imbalance in *Mtb*

We next examined whether AG exposure elicits redox stress in *Mtb*. The down-regulation of aerobic type NADH dehydrogenase I and ATP synthase upon AG exposure may result in reduction of electron flow through respiratory complexes. It might induce back-pressure in respiratory chain causing an increase in cellular NADH/NAD^+^ ratio. We measured NADH/NAD^+^ ratio of *Mtb* exposed to 10X MIC of AG at various time points post-treatment. At pre-lethal stage (6 h post-treatment), we did not observe any change in NADH/NAD^+^ ratios (Fig. 4A). However, a significant elevation of NADH/NAD^+^ ratio was detected 24 h post-treatment, which coincides with AG-induced killing in *Mtb* (Fig. 4A). Our results align with the expression data showing up-regulation of alternate respiratory complexes (*ndh*, *cydAB*, and *nirBD*), ostensibly to replenish oxidized form of NADH. However, prolonged AG exposure results into breakdown of NADH/NAD^+^ balance and killing. Our AG-response network indicates a clear activation of the systems associated with mitigation of oxidative stress in *Mtb*. Therefore, we next determined accumulation of ROS by staining with an oxidant-sensitive fluorescent dye; 2',7'-dichlorofluorescein diacetate (DCFDA) in *Mtb* cells treated with AG (10X MIC) for 3 and 6h. Early time points were considered for ROS measurements to disregard the possibility of death-mediated increase in ROS upon AG treatment. A consistent increase (~ 3-fold increase) in DCFDA fluorescence was observed at both time points as compared to untreated control (Fig. 4B). Under aerobic conditions, ROS is mainly generated through univalent reduction of _O2_ by reduced metals, flavins, and quinones (24), which mainly generates superoxide 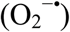. Therefore, we next determined 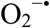 production using a well-established and freely cell-permeable 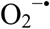 indicator, dihydroethidium (DHE) (25). It is known that DHE specifically reacts with 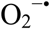 to release fluorescent product 2-hydroxyethidium (2-OH-E^+^), which can be conveniently detected by HPLC (25). The reaction of DHE with other oxidants produces ethidium (E^+^) (55). Owing to biosafety challenges associated with a BSL3 class pathogen such as *Mtb* for HPLC, we measured 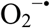 levels inside a related but nonpathogenic strain of mycobacteria, *Mycobacterium bovis* BCG upon AG challenge. BCG cells were treated with AG (10X MIC) for 3 and 6h, followed by DHE staining and HPLC. We found that BCG cells treated with AG generate peaks corresponding to 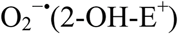 and other ROS (E^+^) (Fig. 4C). The intensity of peaks was significantly higher at 6 h post-treatment as compared to untreated control (Fig. 4C). As a control, we used a well-known 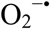 generator (menadione) in our assay and similarly detected a 2-OH-E^+^ peak (Fig. 4C, *inset*). Next, we determined whether thiol-based antioxidant thiourea can reverse the influence of AG on viability of *Mtb*. Thiourea has recently been shown to protect *Mtb* from oxidative stress by modulating the expression of antioxidant genes (37). *Mtb* was co-incubated with various concentrations of thiourea and AG, and viability was measured after 10 days. Thiourea did not exert significant effect on the survival of *Mtb* under normal growing conditions (Fig. 4D), however, it did increase the survival of *Mtb* treated with 0.625X and 1.25X MIC by ~ 10- and 5- fold, respectively (Fig. 4D). At higher AG concentrations (2.5X MIC), only 100 mM of thiourea showed a 2 fold protective effect (Fig. 4D).

**Fig 4:**
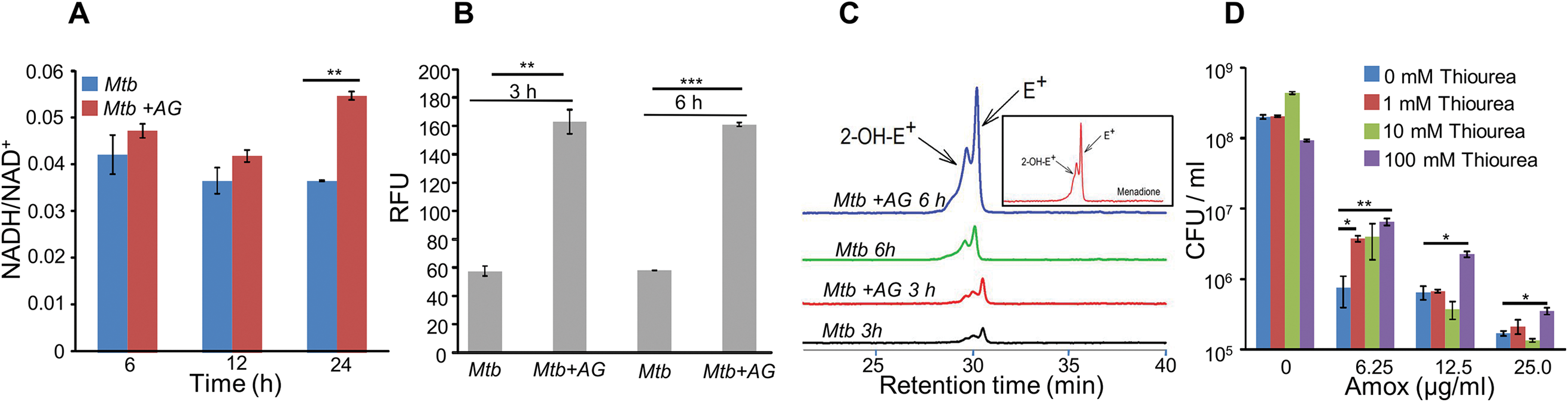
AG influences the internal redox physiology of *Mtb*. Wt *Mtb* or *M. bovis* BCG was grown to an OD_600_ of 0.4 and treated with 100 µg/ml of Amox and 8 µg/ml of Clav (AG). At indicated time points, cells were analyzed for (A) NADH/NAD^+^ estimation, (B) ROS measurement using oxidant-sensitive fluorescent dye; 2',7'-dichlorofluorescein diacetate (DCFDA), and (C) Superoxide estimation using dihydroethidium (DHE) as described in *Materials and Methods*. (D) Wt *Mtb* was grown as describe earlier and exposed to specified concentrations of Amox in presence of 8 µg/ml of Clav for 10 days in presence or absence of thiourea and survival was measured using colony-forming unit (CFU) counts. Error bars represent standard deviations from mean. * p≤0.05, * p≤0.01 and *** p≤0.001. Data are representative of at least two independent experiments done in duplicate.

The above data indicate that bactericidal consequences of AG may be dependent upon internal oxidant-antioxidant balance of *Mtb*. To unambiguously demonstrate this, we exploited a mycobacterial specific non-invasive biosensor (Mrx1-roGFP2) to measure the redox potential of a physiologically-relevant and abundant cytoplasmic antioxidant, MSH (4). Any changes in the oxidation-reduction state of MSH can be reliably quantified by measuring ratio of emission at 510 nm at 405/488 nm excitations (4). *Mtb* expressing Mrx1-roGFP2 was treated with lower (2 μg/ml of Amox+ 4 μg/ml of Clav; 0.2X MIC) and higher (100 μg/ml of Amox+ 8 μg/ml of Clav; 10X MIC) concentrations of AG and intramycobacterial *E_MSH_* was determined by measuring biosensor ratiometric response over time, as described previously (4). To preclude the effect of killing on *E_MSH_* measurements, we tracked biosensor ratios during pre-lethal phase (3 h and 6 h for 10X MIC and 12 h and 24 h for 0.2X MIC) of AG challenge. We observed a modest but consistent increase in 405/488 ratio at 6 h and 24 h post-treatment with 10X and 0.2X MIC of AG, respectively (Fig. 5A), indicating that antioxidant mechanisms are mostly efficient in minimizing the impact of AG-mediated ROS generation on internal *E_MSH_* of *Mtb*.

**Fig 5:**
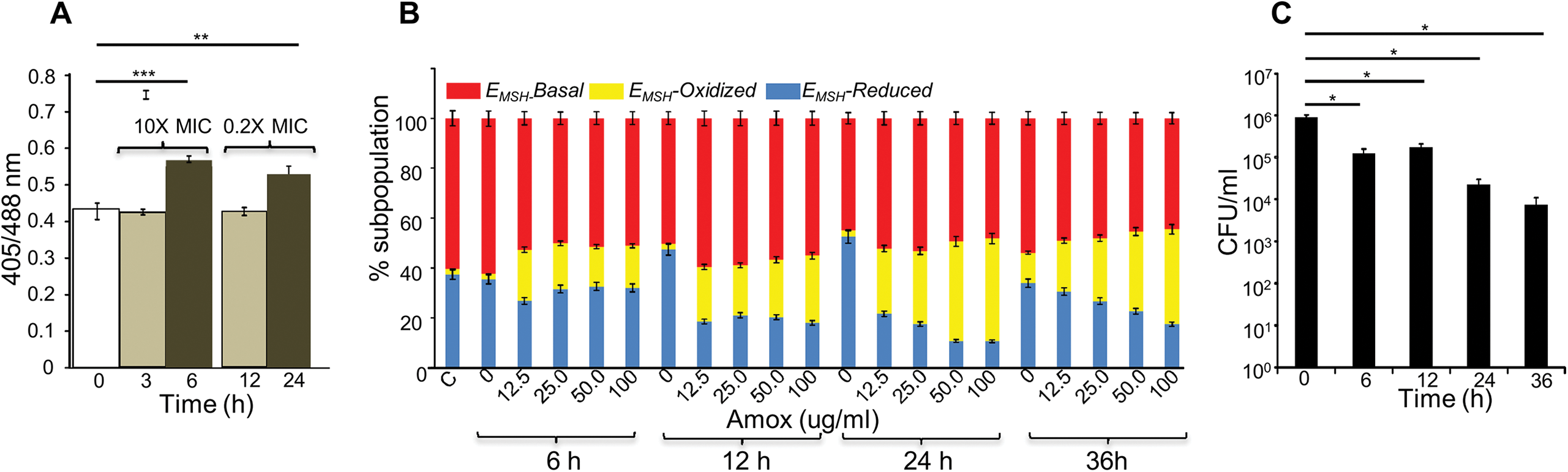
AG induces oxidative shift in *E_MSH_* of *Mtb* in vitro and during infection. (A) Wt *Mtb* expressing Mrx1-roGFP2 was treated with lethal (10X MIC) and sub-lethal (0.2 X MIC) concentrations of AG and ratiometric sensor response was measured at indicated time points by flow cytometry. (B) PMA differentiated THP-1 cells were infected with *Mtb* expressing Mrx1-roGFP2 (moi: 10) and treated with indicated concentrations of Amox in presence of 8 µg/ml of Clav immediately after infection. At indicated time points, ~ 30,000 infected macrophages were analyzed by flow cytometry to quantify changes in *Mtb* subpopulations displaying variable *E_MSH_* as described in *Materials and Methods*. (C) In parallel experiments, infected macrophages were lysed and bacillary load was measured by plating for CFU. Error bars represent standard deviations from mean. * p≤0.05, ** p≤0.01 and *** p≤0.001. Data are representative of at least two independent experiments done in duplicate.

Importantly, to determine whether AG-induced oxidative stress is during infection, we measured dynamic changes in *E_MSH_* of *Mtb* during infection inside human macrophage cell line (THP-1). Infected macrophages were exposed to AG (1.25-fold to 10-fold of the *in vitro* MIC) and the redox response was measured by flow cytometry. As reported earlier, inside macrophages *Mtb* cells displayed variable *E_MSH_*, which can be resolved into *E_MSH_*-basal (-270 mv), *E_MSH_*-oxidized (-240 mV), and *E_MSH_*-reduced (-310 mV) subpopulations (4). Treatment with AG significantly increased oxidized subpopulation over time (Fig. 5B). We observed a time and concentration dependent oxidative shift in *E_MSH_* of *Mtb* subpopulations (Fig. 5B). In parallel, we examined whether the elevated oxidative stress correlates with the killing potential of AG during infection. Macrophages infected with *Mtb* were treated with 10X MIC of AG and bacillary load was monitored by enumerating colony-forming units (CFUs) at various time points post-infection. At 6 h and 12 h post-AG treatment, the effect on *Mtb* survival was marginal (Fig. 5C). However, an ~ 100-fold decline in CFU was observed at 24 h and 36 h post-AG treatment (Fig. 5C). Furthermore, an increase in *E_MSH_*-oxidized subpopulation occurred at a time point where survival was not considerably affected (6 h) (Fig. 5B and 5C), indicating that AG-mediated oxidative stress precedes bacterial death inside macrophages and that the intramycobacterial oxidative stress is not a consequence of AG-induced toxicity. Altogether, data show that AG perturbs mycobacterial redox physiology and the environment inside macrophages potentiates the mycobactericidal effect of AG.

### Mycothiol buffer protects *Mtb* from AG-mediated killing

Since AG induces intramycobacterial oxidative stress, it is likely that the loss of major intracellular antioxidant, mycothiol, might potentiate the antimycobacterial activity of AG. To examine this, we used a MSH negative strain (*MsmΔmshA*) of *Mycobacterium smegmatis (Msm),* an organism that is widely used as a surrogate for pathogenic strains of *Mtb*. Wt *Msm* and *MsmΔmshA* strains were exposed to various concentrations of Amox at a saturating concentration of Clav (8 μg/ml) and percent growth inhibition was measured using Alamar blue (AB) assay. AB is an oxidation-reduction indicator dye that changes its color from non-fluorescent blue to fluorescent pink upon reduction by actively metabolizing cells, whereas inhibition of growth by antimycobacterial compounds interferes with AB reduction and color development (55). As shown in Fig. 6A, at a fixed Clav concentration, *MsmΔmshA* exhibited ~ 3- and 10-fold increased inhibition at 5 μg/ml and 2.5 μg/ml of Amox as compared to wt *Msm*, respectively. At 10 μg/ml of Amox, both strains showed nearly complete inhibition (Fig. 6A). Next, we measured susceptibility to Clav at a fixed concentration of Amox (10 μg/ml). Higher concentrations of Clav (10 μg/ml) inhibited the growth of *Msm* and *MsmΔmshA* with a comparable efficiency (Fig. 6B). However, whereas wt *Msm* overcomes the inhibitory effect of Amox at lower Clav concentrations, *MsmΔmshA* remained sensitive to Amox even at a lowest concentration of Clav (0.625 μg/ml) (Fig. 6B). As shown in figure 6B, *MsmΔmshA* exhibited an ~ 7-fold greater inhibition at 0.625 μg/ml of Clav as compared to wt *Msm*.

**Fig 6:**
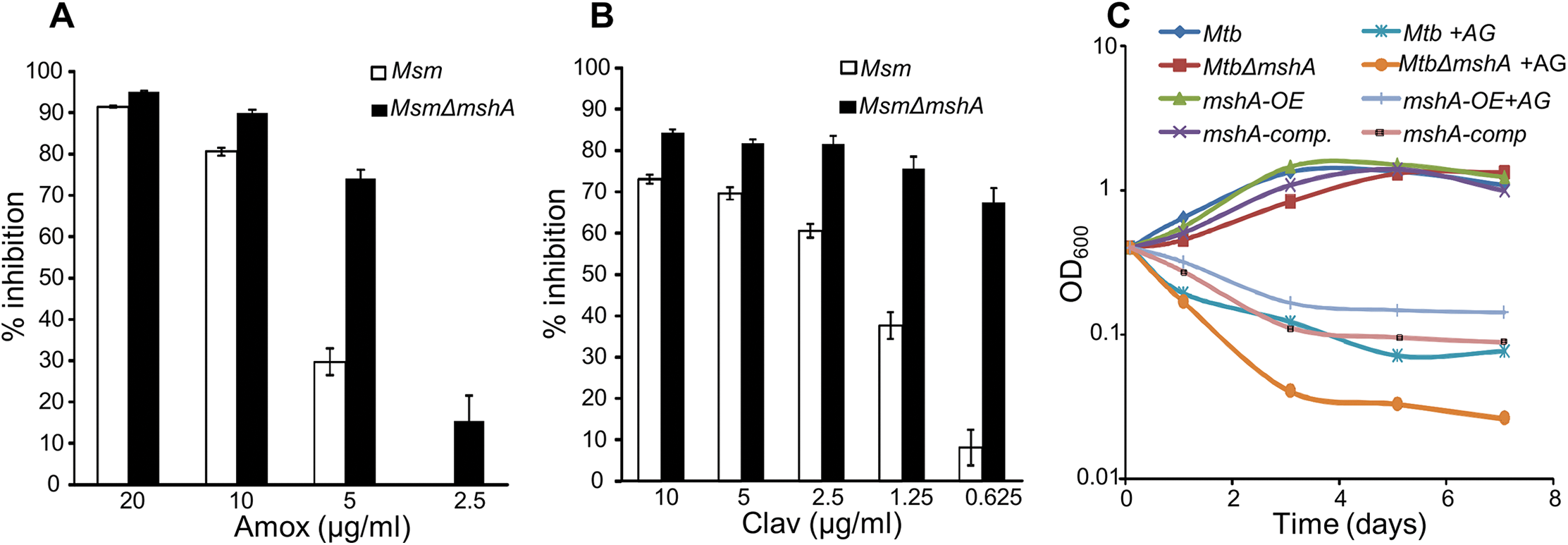
Mycothiol mediates tolerance to AG. Wt *Msm* and *MsmΔmshA* strains were grown to OD_600_ of 0.4 and either treated with various concentrations of (A) Amox at a fixed concentration of Clav (8 µg/ml) or (b) Clav at a fixed concentration of Amox (10 µg/ml) and % inhibition in growth was measured by Alamar blue (AB) assay as described in *Materials and Methods*. (C) Wt *Mtb*, *MtbΔmshA*, *mshA-comp* and *mshA:OE* strains were grown in the presence of 100 µg/ml of Amox and 8 µg/ml of Clav (10X MIC of AG) and survival was monitored by measuring OD_600_ over time. Error bars represent standard deviations from mean. Data are representative of at least two independent experiments done in duplicate.

To confirm that above findings can be recapitulated in slow growing pathogenic mycobacteria (*i.e. Mtb* H37Rv) and to determine the contribution of *E_MSH_* in AG tolerance, we shifted the intramycobacterial *E_MSH_* of *Mtb* towards reductive by conditionally overexpressing *mshA* in *Mtb* (*mshA:OE*) using an anhydrotetracyline (Atc) inducible system (TetRO) (33). We performed *E_MSH_* measurements and confirmed that the overexpression of *mshA* shifted ambient *E_MSH_* of *Mtb* from -275±3 mV to -300±5 mV, indicating an overall elevation in anti-oxidative potential of *Mtb*. The MSH-deficient strain of *Mtb* (*MtbΔmshA*) displayed oxidative *E_MSH_* of >-240 mV, whereas *mshA* complemented strain (*mshA-comp*) displayed *E_MSH_* comparable to wt *Mtb* (*i.e.* -275±3 mV). Wt *Mtb*, *mshA:OE*, *MtbΔmshA* and *mshA-comp* were exposed to 10X MIC of AG (100 μg/ml of Amox+ 8 μg/ml of Clav) and growth was monitored over time by measuring culture density at OD_600_. AG treatments resulted in time dependent decrease in the growth of *Mtb* strains (Fig. 6C). However, the decline was severe in case of *MtbΔmshA* as compared to wt *Mtb*, whereas *mshA-OE* grew relatively better than wt *Mtb* (Fig. 6C). Expression of *mshA* from its native promoter (*mshA-comp*) restored tolerance comparable to wt *Mtb* (Fig. 6C). In sum, AG exposure triggers ROS production and mycothiol provides efficient tolerance towards AG.

### *Mtb* WhiB4 modulates gene expression and maintains *E_MSH_* in response to AG

Altered expression of oxidative-stress genes, elevation of ROS, and perturbation of *E_MSH_* upon AG exposure suggest that intramycobacterial redox potential can serve as an internal cue to monitor the presence of β-lactams. Canonical redox sensors such as OxyR, SoxR, and FNR are either absent or rendered non-functional in *Mtb* (6,9). We have recently shown that *Mtb* features a Fe-S cluster containing transcription factor, WhiB4, which responds to oxidative stress by regulating the expression of antioxidant genes (6). The fact that *whiB4* expression is uniformly repressed by β-lactams (*e.g.,* meropenem and AG) (30) and oxidative stress (6), implicate WhiB4 in linking oxidative stress with β-lactam response in *Mtb*. We directly tested the role of WhiB4 in β-lactam tolerance. We performed microarray of *MtbΔwhiB4* upon treatment with AG (10X of *Mtb* MIC) for 6 h as described previously. A total of 495 genes were induced (≥1.5-fold; *p*-value ≤0.05) and 423 were repressed (≥1.5-fold, *p*-value ≤0.05) in the *MtbΔwhiB4* as compared to wt *Mtb* upon AG-treatment (Supplementary Table S5). Our network analysis showed that diverse functional classes such as cell wall processes, virulence adaptation pathways, intermediary metabolism and respiration, and lipid metabolism were affected in *MtbΔwhiB4* upon AG exposure (Fig. 7A). Microarray data indicated higher expression of genes known to tolerate β-lactams in *MtbΔwhiB4*. Transcription of *blaR* and *blaC* was induced 8.43±4.75 and 2.23±0.19 fold, respectively, in *MtbΔwhiB4* as compared to wt *Mtb* upon treatment (Fig. 7B, Supplementary Table S5). Other genetic determinants of β-lactam tolerance such as PG biosynthetic genes (*murE, murF, and murG*), penicillin-binding proteins (*Rv2864c, Rv3627c and Rv1730c*), and cell division and DNA transaction factors (*ftsK* and *fic*) (Fig. 7B) were up-regulated in *MtbΔwhiB4* upon treatment. *MtbΔwhiB4* also exhibited greater expression of DNA repair genes (SOS response), many of which are known to interfere with cell division and promote β-lactam tolerance in other bacterial species (34) (Fig. 7C). Since transcriptional data implicate WhiB4 in regulating the biosynthesis of PG polymer, we stained PG polymer of wt *Mtb*, *MtbΔwhiB4*, and *whiB4-OE* cells using fluorescent derivative of PG binding antibiotic, vancomycin (Bodipy-VAN) and imaged cells under confocal microscope. The *whiB4-OE* strain harbors pEXCF*-whiB4* plasmid, which allows overproduction WhiB4 in the presence of an inducer anhydrotetracycline (ATc) (6). As expected, poles of *wt Mtb* and *whiB4-OE* cells were fluorescently labeled, consistent with the incorporation of nascent PG at the poles in mycobacteria (Supplementary Fig. S4A) (53). Interestingly, Van-FL was found to label entire length of *MtbΔwhiB4*, indicating deposition of PG along the entire body of the cylindrical cells (Supplementary Fig. S4A). *MtbΔwhiB4* cells were also marginally longer than wt *Mtb* (Supplementary Fig. S4B). Further experimentations are required to comprehensively understand how WhiB4 modulates PG biosynthesis. Nonetheless, our transcriptomics and imaging data are in reasonable agreement with each other and support the PG-regulatory function of WhiB4 in *Mtb*.

**Fig 7:**
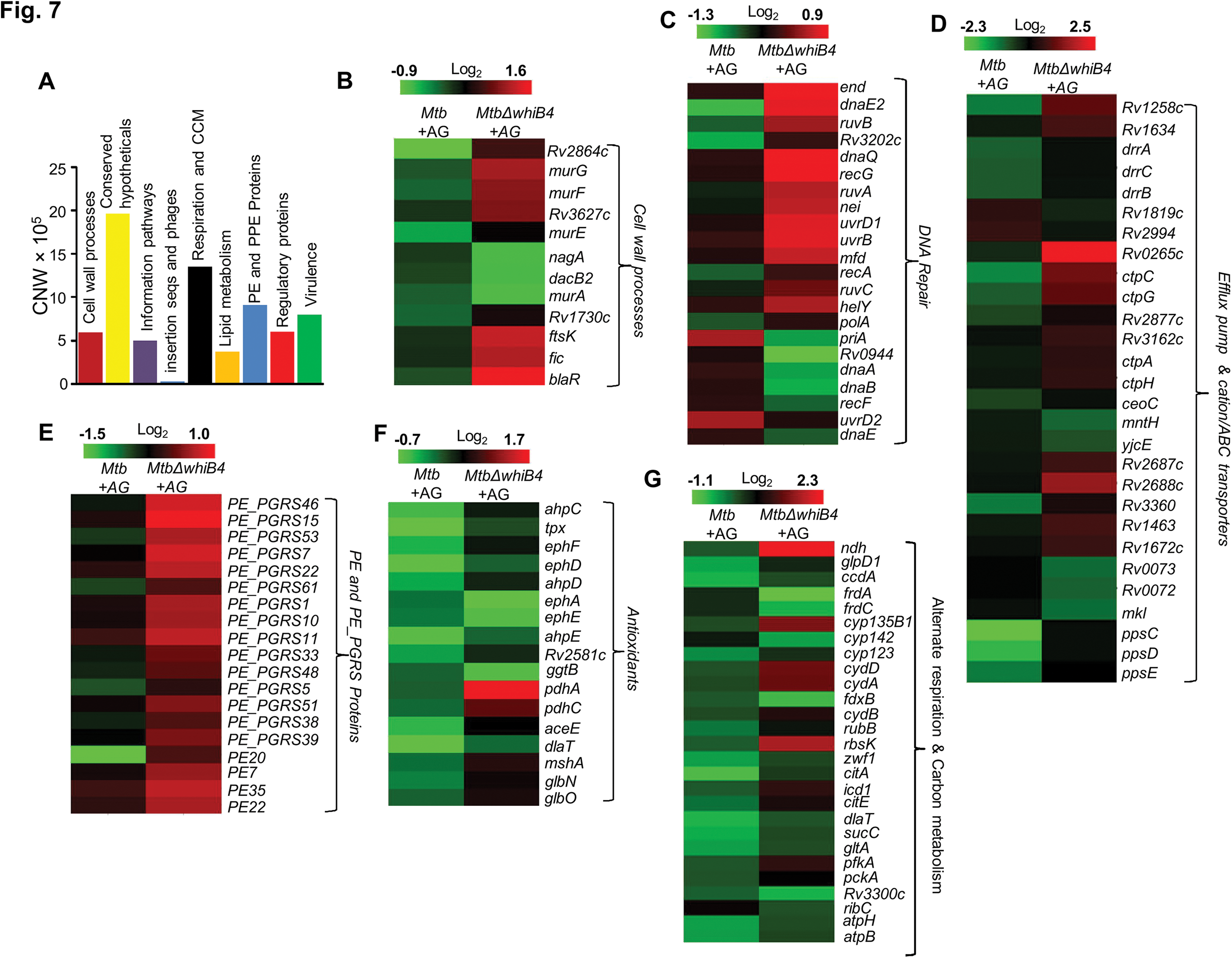
WhiB4 regulates response to AG in *Mtb*. (A) Cumulative node weight intensities (CNW) of different functional classes regulated by WhiB4 upon AG treatment. (B-G) Heat maps depicting fold-change (log_2_) in expression of genes coordinating cell wall processes, alternate respiration and central metabolism, antioxidants, DNA repair, PE and PE_PGRS and drug efflux pumps in case of *Mtb* and *MtbΔwhiB4* treated with AG for 6 h as described in *Materials and Methods*.

Other mechanisms that could link WhiB4 with drug tolerance include the heightened expression of cation transporters, ABC-transporters, PDIM lipid biogenesis (*ppsC, ppsD, ppsE, drrA, drrB,* and *drrC*) and drug-efflux pumps (*Rv1258c* and *Rv1634*) in *MtbΔwhiB4* upon AG exposure (Fig. 7D). Lack of Rv1258c pump and PDIM lipids have been reported to sensitize *Mtb* towards β-lactams and vancomycin (10,51). Several members of PE_PGRS genes involved in maintaining cell wall architecture and protection from oxidative stresses were up-regulated in *MtbΔwhiB4* (Fig. 7E) (13). Our results also revealed that redox-metabolism is significantly altered in *MtbΔwhiB4* in response to AG. For example, components of the NADH-dependent peroxidase (*ahpCD*), peroxynitrite reductase complex (*dlaT*), thiol-peroxidase (*tpx*), mycothiol biosynthesis (*mshA*), pyruvate dehydrogenase complex (*pdhA, pdhC,* and *aceE*), and hemoglobin like proteins (*glbO*) were induced in AG-challenged *MtbΔwhiB4* compared to wt *Mtb* (Fig. 7F). Importantly, most of these enzymatic activities are well known to confer protection against oxidative and nitrosative stress in *Mtb* (21,31,32,40,56,58). Further, similar to wt *Mtb*, primary NADH dehydrogenase complex (*nuo operon*) was down-regulated in *MtbΔwhiB4* in response to AG treatment (Supplementary table S6). However, compensatory increase in alternate respiratory complexes such as *ndh* and *cydAB* was notably higher in *MtbΔwhiB4* than wt *Mtb*, indicating that *MtbΔwhiB4* is better fit to replenish reducing equivalents during drug-induced cellular stress (Fig. 7G). In line with this, components of TCA cycle and pentose phosphate pathway involved in generating cellular reductants (NADH and NADPH) were induced in *MtbΔwhiB4* as compared to wt *Mtb*.

Overall AG-exposure elicits transcriptional changes, which are indicative of a higher potential of *MtbΔwhiB4* to maintain redox homeostasis upon drug exposure. We directly assessed this by examining changes in *E_MSH_* of *MtbΔwhiB4* in response to AG *in vitro* and inside macrophages. Under both culture conditions, *MtbΔwhiB4* robustly maintained intramycobacterial *E_MSH_*, whereas overexpression of *whiB4* in *MtbΔwhiB4* showed a significant oxidative shift (Supplementary Fig. S5). In sum, our results suggest that WhiB4 can mediate AG tolerance by regulating multiple mechanisms including PG biogenesis, SOS response, and antioxidants.

### *Mtb* WhiB4 regulates BlaC in a redox-dependent manner

Data indicate that WhiB4 modulates the expression of genes involved in β-lactam tolerance (*blaR* and *blaC*) and redox metabolism. Using qRT-PCR, we confirmed that the expression of *blaR* and *blaC* upon AG exposure was 8.43±4.75 and 2.23±0.19 fold higher, respectively, in *MtbΔwhiB4* as compared to wt *Mtb*. Next, we examined WhiB4 interaction with the *blaC* promoter region using EMSA. Previously, we have shown that WhiB4 lacking Fe-S cluster (apo-WhiB4) binds DNA upon oxidation of its cysteine thiols, while reduction abolished DNA binding (6). We generated thiol-reduced and – oxidized forms of apo-WhiB4 as described previously (6). The oxidized and reduced apo-WhiB4 fractions were incubated with ^32^P-labeled promoter fragments (~150 bp upstream) of *blaC and blaR* and complex formation was visualized using EMSA.

As shown in figure 8A and 8B, oxidized apo-WhiB4 binds DNA in a concentration-dependent manner, whereas this binding was significantly reversed in case of reduced apo-WhiB4. Since WhiB4 bind to its own promoter (6), we confirmed that oxidized apo-WhiB4 binds to its promoter in concentrations comparable to that required for binding *blaC* and *blaR* promoters (Fig. 8C). Next, we performed *in vitro* transcription assays using a highly sensitive *Msm* RNA polymerase holoenzyme containing stoichiometric concentrations of principal Sigma factor, SigA (RNAP-σA) (6) and determined the consequence of WhiB4 on *blaC* transcript. As shown in figure 8D, addition of oxidized apo-WhiB4 noticeably inhibited transcription from *blaC* promoter, whereas reduced apo-WhiB4 restored normal levels of *blaC* transcript. Lastly, we directly measured BlaC activity in the cell-free extracts derived from wt *Mtb*, *MtbΔwhiB4*, and *whiB4-OE* strains using a chromogenic β-lactam nitrocefin as a substrate (15). Using *in vivo* thiol-trapping assay, we have earlier shown that WhiB4 predominantly exists in an oxidized apo-form upon overexpression in *whiB4-OE* strain (6). Therefore, *whiB4-OE* strain will reveal the role of oxidized apo-WhiB4 in BlaC regulation *in vivo*. Cell free extracts of *MtbΔwhiB4* possessed ~70% higher and *whiB4-OE* showed ~30% reduced nitrocefin hydrolysis as compared to wt *Mtb*, respectively (Fig. 8E). Decreased BlaC activity upon WhiB4 overexpression is in agreement with oxidized apo-WhiB4-mediated repression of the *blaC* transcription *in vitro*. To clarify the physiological relevance of redox- and *whiB4*- dependent transcription of *blaC*, we shifted the internal redox balance of *whiB4-OE* using a cell permeable thiol-oxidant, diamide (5 mM), or a thiol-reductant, DTT (5 mM), and measured nitrocefin hydrolysis by cell free extracts. We have previously reported that treatment with 5 mM diamide or DTT did not adversely affect growth of *Mtb* (50). Pretreatment of *whiB4-OE* with DTT largely restored BlaC activity to *MtbΔwhiB4* levels, whereas diamide did not lead to further decrease in BlaC activity in *whiB4-OE* strain (Fig. 8F). Effective reduction of oxidized apo-WhiB4 by DTT within *whiB4-OE* cells may have led to loss of WhiB4 DNA binding and transcription repressor activity, thereby causing elevated *blaC* expression and activity. Taken together, these results led us to conclude that WhiB4 regulates β-lactamase expression and activity in a redox-dependent manner.

**Fig 8:**
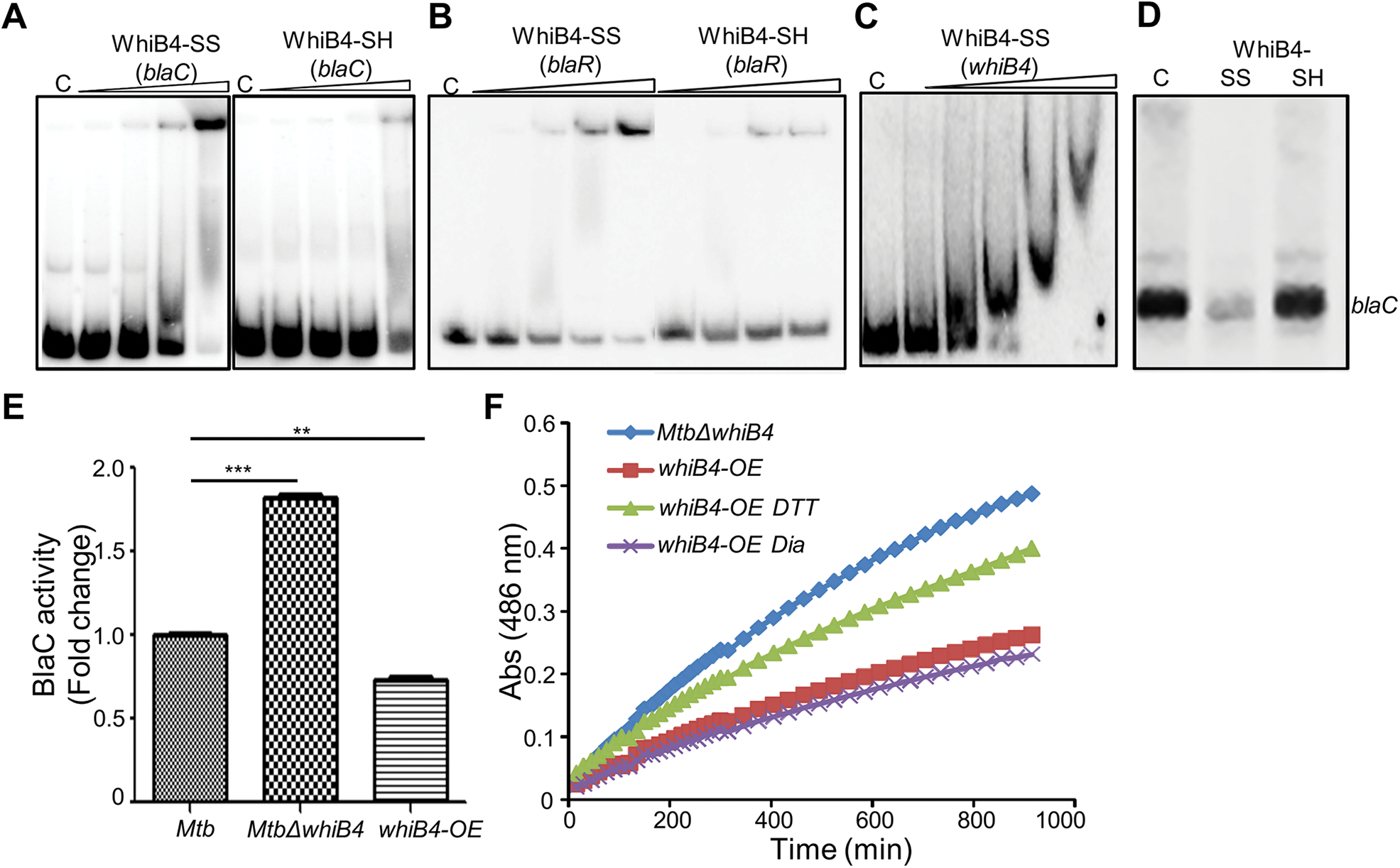
Regulation of β-lactamase by WhiB4 in a redox-dependent manner. Oxidized (WhiB4-SS) and reduced (WhiB4-SH) forms of apo-WhiB4 were prepared. The concentrations of apo-WhiB4 used for EMSAs were 0.5, 1, 2, and 4 μM. EMSA reactions were performed with 0.5 nM ^32^P-labelled *blaC* (A), *blaR* (B) and *whiB4* (C) promoter DNA fragments. C: DNA binding in the absence of WhiB4 in each panel. (D) Effect of WhiB4 on *in vitro* transcription. Single round transcription assays show that RNAP-σ^A^ efficiently directs transcription from *blaC* promoter. 100 nM of *blaC* promoter DNA fragment was pre-incubated with either 1 μM WhiB4-SS or WhiB4-SH and subjected to transcription by RNAP-σ^A^ as described in *Materials and Methods*. C: *blaC* transcript in the absence of WhiB4. (E) 100 μg of cell free lysates derived from exponentially grown (OD_600_ of 0.6) wt *Mtb*, *MtbΔwhiB4* and *whiB4-OE* were used to hydrolyze nitrocefin. β-lactamase activity was measured by monitoring absorbance of hydrolyzed nitrocefin at 486 nm as described in *Materials and Methods*. The fold change ratios clearly indicate a significantly higher and lower β-lactamase activity in *MtbΔwhiB4* and *whiB4-OE*, respectively, as compared to wt *Mtb. P*-values are shown for each comparison. (F) *whiB4-OE* strain was pre-treated with 5 mM DTT or Diamide and β-lactamase activity in cell free lysates was compared to *MtbΔwhiB4* over time. * p≤0.05, ** p≤0.01 and *** p≤0.001. Data are representative of at least two independent experiments done in duplicate.

### WhiB4 regulates survival in response to β-lactams in *Mtb*

Based on above results, we hypothesize that WhiB4 -sufficient and –deficient strains would have differential susceptibility towards β-lactams. We found that *MtbΔwhiB4* uniformly displayed ~ 4-8 folds higher MICs against β-lactams as compared to wt *Mtb* (Table 1). This effect was specific to β-lactams, as the loss of WhiB4 did not alter MICs for other anti-TB drugs such as INH and RIF (Table 1). More-interestingly, over-expression of WhiB4 displayed ~ 2-4 folds greater sensitivity towards β-lactams as compared to wt *Mtb* (Table 1). We predicted that if WhiB4 is controlling tolerance to β-lactams by regulating *blaC* expression, we would see variations in inhibitory concentrations of Clav against wt *Mtb*, *MtbΔwhiB4,* and *whiB4-OE* (in presence of a fixed concentration of Amox). As expected, inhibition of *MtbΔwhiB4* by 10 μg/ml of Amox requires 4-fold and 8-fold higher Clav as compared to wt *Mtb* and *whiB4-OE* strains, respectively (Fig. 9A). Phenotypic data is in complete agreement with the higher and lower BlaC activity in *MtbΔwhiB4* and *whiB4-OE*, respectively. Studies in animals and humans have demonstrated higher efficacy of β-lactams and β-lactamase inhibitor combination against MDR/XDR-TB. Our results show that WhiB4 overexpression significantly elevated the capacity of β-lactams to inhibit drug-sensitive *Mtb*. To investigate whether WhiB4 overexpression similarly affects growth of drug-resistant strains, we over-expressed WhiB4 in clinical strains isolated from Indian patients (single drug resistant [SDR; BND320], multi-drug resistant [MDR; Jal 2261 and Jal 1934] and extensively drug-resistant [XDR; MYC 431]) (4) and determined sensitivity towards 10 μg/ml of Amox at various concentrations of Clav. Consistently, over-expression of WhiB4 uniformly resulted in an ~2-4 –fold increased sensitivity of drug-resistant strains towards Amox and Clav combinations (Fig. 9B, 9C, 9D, 9E).

**Table 1.** Minimum Inhibitory Concentrations (MICs) of Cell wall targeting drugs.

| *Drugs* | *μg/ml* |  |  |
| --- | --- | --- | --- |
|  | *Mtb* | *MtbΔWhiB4* | *whiB4-OE* |
| Amoxicillin | 80 | >160 | 40 |
| Ampicillin | 500 | 4000 | 250 |
| Cloxacillin | 400 | 800 | 200 |
| Carbenicillin | 1024 | 4096 | 512 |
| Meropenem | 5 | 20 | 2.5 |
| Penicillin | 200 | 800 | 100 |
| Lysozyme | 50 | 200 | 25 |
| Vancomycin | 10 | 80 | 2.5 |
| Isoniazid | 0.0625 | 0.0625 | 0.03125 |
| Rifampicin | 0.0625 | 0.0625 | 0.0625 |

**Fig 9:**
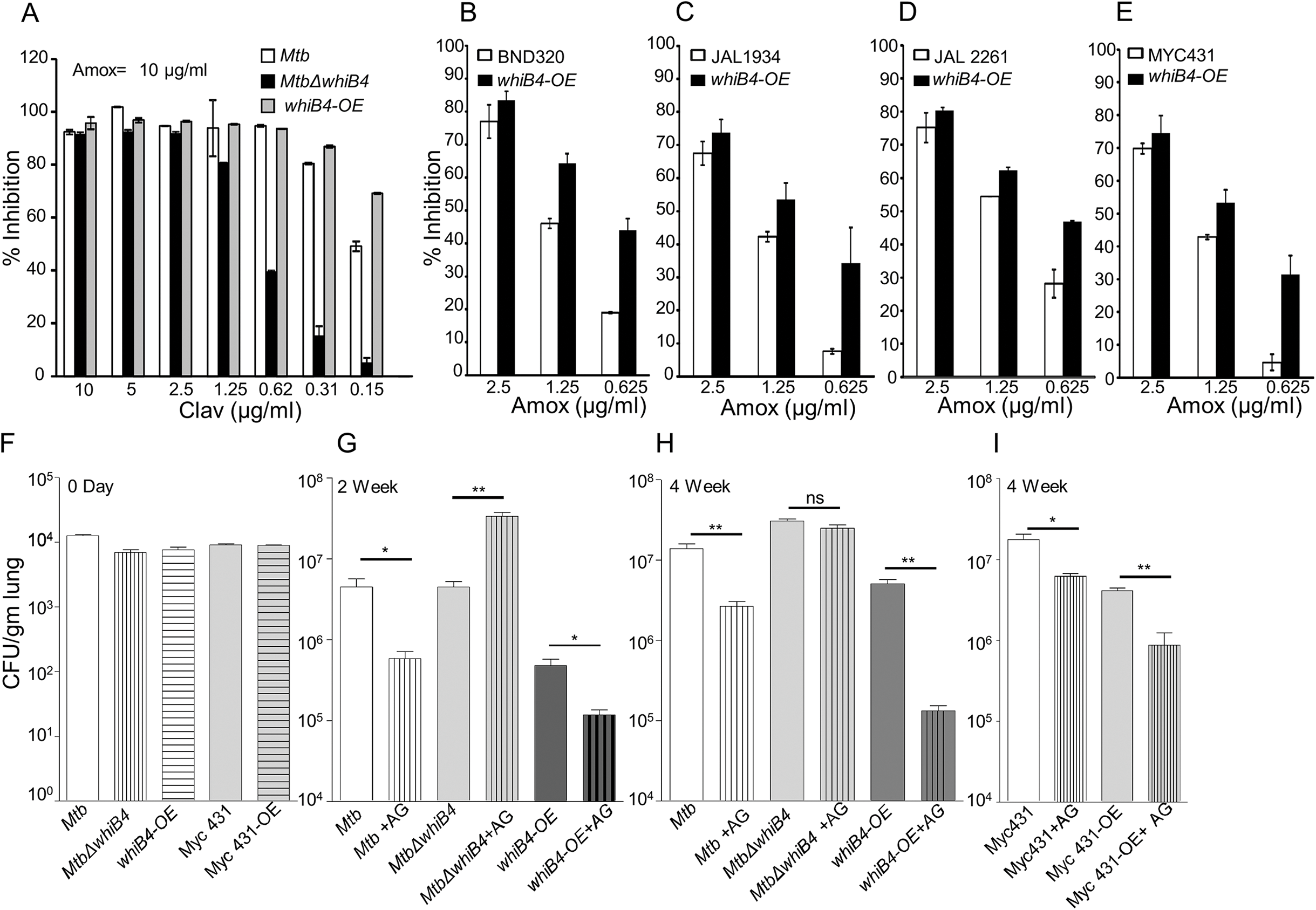
WhiB4 regulates AG tolerance in drug-sensitive and –resistant strains of *Mtb*. (A) Wt *Mtb*, *MtbΔwhiB4* and *whiB4-OE* were incubated with different concentrations of Clav at a fixed concentration of Amox (10 µg/ml) and % inhibition of growth was measured by AB assay as described in *Materials and Methods*. To determine if WhiB4 modulates the sensitivity of AG in drug-resistant strains, WhiB4 was over-expressed in clinical strains (B) BND 320 (C) Jal 1934, (D) Jal-2261, and (E) MYC 431 and cells were incubated with different concentrations of Amox at a fixed concentrations of Clav (8 µg/ml). The percent growth inhibition was measured by AB assay as described in *Materials and methods*. WhiB4 modulates susceptibility to AG during acute infection in mice (F-I). Inbred BALB/c mice (n=3) given an aerosol challenge of various *Mtb* strains and orally administered with Amox (200 mg/kg of body weight) and Clav (50 mg/kg of body weight) twice a day. Survival of *Mtb* strains was assessed for bacterial burden in lungs by CFU analysis. Statistical significance for the pulmonic bacterial load was obtained by comparing CFU obtained from untreated animals (vehicle control) as compared to drugs treated animals infected with various *Mtb* strains. * p≤0.05, ** p≤0.01 and *** p≤0.001.

Lastly, we asked if WhiB4 influences tolerance to AG during infection. Poor half-life of AG in mice makes it challenging to assess the efficacy of AG *in vivo* (43). However, AG induces a marginal (~0.5 log reduction) killing of *Mtb* in an acute model for TB infection in mice (52). Therefore, we compared bacillary load of *Mtb* strains in the lungs of mice during acute infection (see *Materials and Methods*). Approximately 10^4^ bacterial cells were implanted in lungs of BALB/c mice (Fig. 9F) and at 3-days post-infection mice were treated with AG (200 mg/kg body weight of Amox and 50 mg/kg body weight of Clav) twice a day for 2 and 4 –weeks. Bacterial numbers were determined in the infected lungs treatment. At 24h post-infection, bacillary load was comparable between *Mtb* strains (Fig. 9F). At 2 and 4 weeks post-treatment, wt *Mtb* exhibited ~7-fold reduction in bacillary load than untreated mice (Fig. 8G and 8H). Overexpression of WhiB4 resulted in ~4- and ~38-fold decline in CFU at 2 and 4 weeks, post-treatment, as compared to untreated animals (8G and 8H). In contrast, *MtbΔwhiB4* either displayed an increase (~8-fold) or maintained comparable bacillary load at 2 and 4 weeks post-treatment, respectively, relative to untreated mice (Fig. 8F, 8G, and 8H). Lastly, we overexpressed WhiB4 in MYC 431 XDR strain and examined AG efficacy in mice as described earlier. Expectedly, WhiB4 overexpression increased the sensitivity of MYC 431 towards AG treatment at 4 weeks post-infection (Fig. 9I). In conclusion, we observed that WhiB4 overexpression significantly attenuates *Mtb* survival *in vivo*, a phenotype most likely due to antioxidant and β-lactamase repressor function of WhiB4.

## Discussion

We identified internal *E_MSH_* of *Mtb* as a crucial determinant of mycobacterial sensitivity to AG, and identified a central role of WhiB4 in regulating gene expression, maintaining redox balance and survival in this process. Down-regulation of TCA cycle genes and up-regulation of glyoxylate cycle in response to AG are consistent with the reports of elevated tolerance to diverse bactericidal antibiotics, including β-lactams, in bacteria with diminished fluxes through TCA cycle (26,38). Aligning with this, metabolomic profiling of *Mtb* in response to other anti-TB drugs elegantly demonstrated that tolerance is accompanied with reduced TCA cycle activity and elevated fluxes through glyoxylate shunt (37). *Mtb* exhibits tolerance to antibiotics during non-replicating persistence in hypoxia (3). Under these conditions, drug tolerance was accompanied by a redirection of respiration from energetically efficient route (*e.g.,* NADH dehydrogenase I) to less energy efficient course (*e.g.,* NDH, CydAB oxidase), and any interference with this respiratory-switch over (*e.g.,* CydAB mutation) led to resensitization of mycobacteria to antibiotics (29,41). This seems to be a unifying theme underlying tolerance to conventional as well as the newly discovered anti-TB drugs bedaquiline (BDQ) and Q203 (27). Agreeing to this, we found that exposure of *Mtb* to AG elicited a transcriptional signature indicating a shift from energy efficient respiration to energetically less favored pathways as shown by a significant induction of *ndh* and *cydAB* transcripts and a down-regulation of *nuo*, *cydbc1*, and *atp A-H*. In bacteria, including *Mtb*, cytochrome bd oxidase also displays catalase and/or quinol oxidase activity (1,29), which confers protection against oxidative stress and nitrosative stress. On this basis, upregulation of cytochrome bd oxidase in response to AG is indicative of oxidative stress in *Mtb*. Bactericidal antibiotics, including β-lactams, have been consistently shown to produce ROS as a maladaptive consequence of primary drug-target interaction on TCA cycle and respiration (11,26,28). While this proposal has been repeatedly questioned, it is strongly reinforced by multiple independent studies demonstrating that tolerance to antibiotics is linked to the bacterial ability to detoxify antibiotic-triggered ROS generation (19,38,49,60). We confirmed that AG stimulates oxidative stress in *Mtb in vitro* and during infection. However, in contrast to other studies (26), oxidative stress was not associated with a breakdown of NADH/NAD^+^ homeostasis, likely reflecting efficient ETC fluxes through NDH and cytochrome bd oxidase. In *Mtb*, rerouting of electron flux through cytochrome bd oxidase increases oxygen consumption (27), which can trigger _O2_^−•^ and H_2O2_ generation by univalent reduction of _O2_ by the metal, flavin, and quinone containing cofactors of the respiratory enzymes (23,24). Recently, it has been shown that intramycobacterial antioxidant buffer, MSH, protects *Mtb* from small molecule endogenous superoxide generators and ROS-generated by vitamin C (55,59). Specific to AG, we found that anti-mycobactericidal activity is greatly potentiated in MSH deficient mycobacterial strains, whereas overexpressing strain displayed tolerance. This is all consistent with the generation of ROS and MSH as key regulatory mechanisms underlying AG tolerance.

Studies indicated the importance of a broader range of physiological programs such as altered metabolic state and oxidative stress as contributory factors in antibiotic resistance. However, it is not clear if specific regulatory mediators exist which can assess physiological changes to regulate both primary drug targets and secondary consequences of primary drug-target interactions to functionally coordinate tolerance. Mechanisms of drug tolerance are either controlled by global changes in bacterial physiology by ppGpp or toxin-antitoxin (TA) modules (20). Further, regulatory systems such as SoxRS in other bacteria and WhiB7 in *Mtb* facilitate physiological changes required for formation of drug-tolerant persisters without specifically affecting the expression of direct targets of antibiotics (2,36). We, for the first time, identified WhiB4 as a transcriptional regulator of both the genetic determinants of β-lactam resistance (*e.g.,* β-lactamase) and physiological changes associated with phenotypic drug tolerance in *Mtb* (*e.g.,* redox balance).

Due to lack of an extracellular β-lactam sensing domain in *Mtb* BlaR, how *Mtb* responds to β-lactam is unknown. While several possibilities including the involvement of serine/threonine protein kinases (PknA/PknB) containing β-lactam interacting PASTA domains are suggested to regulate BlaR-BlaI activity (35,45), our findings implicate internal redox balance and WhiB4 in responding to β-lactams. We detected that oxidized apo-WhiB4 binds and represses the expression of BlaR and BlaC, whereas reduction reversed this effect. Loss of WhiB4 derepresses BlaR and stimulates expression and activity of BlaC, possibility via BlaR-mediated cleavage of the repressor of *blaC* (*i.e.* BlaI). In addition to *blaC*, BlaI also binds to the promoter region of genes encoding cytochrome bd oxidase and ATP synthase (45), both of which showed higher expression in *MtbΔwhiB4*. Altogether it indicates that regulatory function of BlaI is dependent upon the ability of WhiB4 to coordinate *blaR* expression in response to redox changes associated with β-lactam exposure. Our findings indicate a possible regulatory loop between electron transport chain and β-lactam induced oxidative stress where WhiB4/BlaI/BlaR may act as an important link between them. Under unstressed conditions, uncontrolled expression of genes such as *blaC* and *cydAB* is prevented by WhiB4-mediated DNA binding and repression of *blaR-blaI* locus. WhiB4 Fe-S cluster is uniquely sensitive to oxygen and a fraction of WhiB4 exists in the apo-oxidized form inside aerobically growing *Mtb* (6). Since oxidized apo-WhiB4 is known to repress its own expression (6), *Mtb* can down-regulate the expression of *whiB4* by elevating the levels of oxidized apo-WhiB4 in response to oxidative stress caused by β-lactams. The down-regulation of WhiB4 can reduce its negative influence on gene expression, necessary to adjust the expression of *blaI, blaR*, *blaC*, and genes involved in maintaining respiration and redox balance to neutralize β-lactam toxicity. However, considering that WhiB4 binds DNA non-specifically with a preference for GC rich sequences (6), the exact molecular mechanism of how WhiB4 regulates expression in response to β-lactams will be investigated further.

## Innovation

Our study discovered a new redox based mechanism of AG tolerance in *Mtb*. Based on this work, we predict that compounds/drugs targeting bacterial systems that remediate oxidative damage (*e.g.,* 4-butyl-4-hydroxy-1- (4-hydroxyphenyl)-2-phenylpyrazolidine-3,5-dione) (16), elevate endogenous ROS (*e.g.,* clofazimine/vitamin C) (4,59), inhibit respiration (*e.g.,* Q2O3) (27), and block ATP homeostasis (*e.g.,* bedaquiline) (27) can be effective companions in potentiating the action of β-lactam and β-lactamase combinations in *Mtb*.

## Materials and Methods

### Bacterial strains, mammalian cells and growth conditions

The mycobacterial strains were grown aerobically in 7H9 broth (Difco) or 7H11 agar (Difco) supplemented with 0.2% glycerol, Middlebrook Oleic acid Albumin Dextrose-Catalase (OADC) or 1X Albumin Dextrose Saline (ADS) enrichment and 0.1% Tween 80 (broth). *E. coli* cultures were grown in LB medium. Antibiotics were added as described earlier (6). For WhiB4 overexpression, *whib4-OE* strain was grown aerobically to an OD_600_ nm =0.3, followed by induction with 200 ng/ml Anhydro Tetracycline (Atc, Cayman Chemicals) at 37°C for 18 hr. The human monocytic cell line THP-1 was differentiated using 10-15 ng/ml phorbol 12-myristate 13-acetate (PMA; Sigma-Aldrich Co., St. Louis, MO, USA) and cultivated for infection experiments as described previously (39).

### Drug sensitivity assay

Sensitivity to various drugs was determined using microplate alamar blue assay (AB). AB assay was performed in 96 well flat bottom plates. *Mtb* or *Msm* strains were cultured in 7H9-ADS medium and grown till exponential phase (OD_600_ ~0.6). Approximately 1×10^5^ bacteria were taken per well in a total volume of 200 µl of 7H9-ADS medium. Wells containing no *Mtb* were the autofluorescence control. Additional controls consisted of wells containing cells and medium only. Plates were incubated for 5 days (*Mtb*) or 16 h (*Msm*) at 37ºC, 30µl (0.02% wt/vol stock solution) Alamar blue (SIGMA-R7017) was added. Plates were re-incubated for color transformation (blue to pink). Fluorescence intensity was measured in a SpectraMax M3 plate reader (Molecular Device) in top-reading mode with excitation at 530 nm and emission at 590 nm. Percentage inhibition was calculated based on the relative fluorescence units and the minimum concentration that resulted in at least 90% inhibition was identified as MIC.

### Intracellular superoxide detection

*Mycobacterium bovis BCG* was cultured in 5 mL of Middlebrook 7H9 medium with 10% albumin-dextrose-saline (ADS) supplement at 37ºC and grown till OD_600_ nm of 0.4. The cultured bacteria were centrifuged to aspirate out the medium and re-suspended with fresh 7H9 medium. This bacterial solution was incubated with AG for 3 h and 6 h time points and 100 μM DHE was added for 1 h in dark by covering the falcon tube in an aluminum foil. The suspension was centrifuged to aspirate out any excess of the compound and DHE in the medium. The collected bacterial pellet was re-suspended with acetonitrile and the cells were lysed using a probe sonicator for 3 min on ice. The cell lysate was then removed by centrifugation and the supernatant was separated and injected in Agilent high performance liquid chromatograph (HPLC) attached with a fluorescence detector (excitation at 356 nm; emission at 590 nm) for analysis. Zorbax SB C-18 reversed-phase column (250×4.6 mm, 5 μm) was used and water: acetonitrile (0.1% trifluoroacetic acid) was applied as mobile phase while flow rate was maintained at 0.5 ml/min. The HPLC method used was as described previously (25).

### Intracellular NADH/ NAD+ ratio

NADH/NAD^+^ ratios upon AG treatment were determined by NAD/NADH Quantification Kit (Sigma-Aldrich, St. Louis, MO, USA). *Mtb* cells were cultured to OD_600_ of 0.4 and treated with Amox-Clav combination for various time points (6,12 and 24 h). 10 ml of culture was harvested and washed with 1X PBS and NADH/NAD^+^ ratio was determined according to manufacturer’s instructions.

### Detection of intracellular ROS

ROS generation upon AG treatment was assessed by using a peroxide detection agent, 5-(and 6)-chloromethyl-2′,7′-dichlorodihydrofluorescein diacetate, acetyl ester (CM-H2DCFDA; Invitrogen). The reagent is converted to a fluorescent product by cellular peroxides/ROS as determined by flow cytometry. *Mtb* cells were cultured to mid-logarithmic phase (OD_600_ of 0.4) and AG treatment was given for 3h and 6 h. At each time point, 500 μL of culture was aliquoted and incubated with 20 μM of CM-H2DCFDA in dark (30 min) at 37°C. Cells were washed with 1X PBS and analyzed by FACS Verse flow cytometer (BD Biosciences, San Jose, CA, USA). CM-H2DCFDA fluorescence was determined (excitation at 488-nm and emission at 530 nm) by measuring 10,000 events/sample.

### Nitrocefin hydrolysis assay

β-lactamase activity in *Mtb* strains was determined spectrophotometrically by hydrolysis of nitrocefin, a chromogenic cephalosporin substrate that contains a β-lactam ring. Bacterial cultures were grown to an OD_600_ of 0.6-0.8 and cells were harvested and lysed using bead beater (FastPrep^®^ Instrument, MP Bio). The cell free lysate was clarified by centrifugation and 100 μg of lysate was incubated with 100 µM Nitrocefin (484400-MERCK). Hydrolysis of nitrocefin was monitored at 486 nm emission (excitation at 390 nm) using SpectraMax M3 plate reader (Molecular Devices) at regular intervals. Fold activity was calculated based on changes in absorbance at 486 nm over time. Normalization was performed by Bradford estimation of total protein in the cell free lysates.

### Microarray hybridisation and data analysis

For microarray analyses, the wt *Mtb* and *MtbΔwhiB4* strains were cultured to an OD_600_ 0.4 and exposed to AG (100 µg/ml of Amox and 8 µg/ml of Clav) for 6 h. For CHP stress, wt *Mtb* grown similarly was treated with 250 µM of CHP for 1 h and samples was processed for microarrays. Total RNA was isolated from samples (taken in replicates), processed and hybridized to *Mtb* Whole Genome Gene Expression Profiling microarray- G2509F (AMADID: G2509F_034585, Agilent Technologies PLC) and data was analysed as described (33). DNA microarrays were provided by the University of Delhi, South Campus, MicroArray Centre (UDSMAC). RNA amplification, cDNA labeling, microarray hybridization, scanning, and data analysis were performed at the UDSMAC as described (33). Slides were scanned on a microarray scanner (Agilent Technologies) and analyzed using GeneSpring software. Results were analyzed in MeV with significance analysis of microarrays considered significant at *p*≤0.05. The normalized data from the microarray gene expression experiment have been submitted to the NCBI Gene Expression Omnibus and can be queried via Gene Expression Omnibus series accession number GSE93091 (AG exposure) and GSE73877 (CHP exposure).

### Constructing of condition-specific networks

Global PPI network was generated using the dataset described in the studies outlined in Supplementary Table S7. After constructing global PPI network of *Mtb*, we then extracted those interactions that are specific for genes present in our transcriptome data. Our microarray specific network consists of 34035 edges and 4016 nodes. The expression data was used for assigning weights to nodes and edge in the PPI network to make it condition-specific. The formalism of node and edge weight calculation is given below

Node weight: We calculated node weight (NW) values for each node in the network by multiplying the normalized intensity values with the corresponding fold-change (FC) values. These values were uniformly scaled by multiplying with 10^4^.

NWi = FCi x Normalized signal intensity

where i denotes the node in the network.

Edge weight: In order to calculate the edge weight values, we first calculated Edge-betweenness (EB) using NetworkX, a python package (https://networkx.github.io/). These values were scaled by multiplying with 10^6^. The node weight values were used to calculate the edge weight (EW) values as follows: -
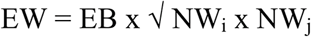
where i and j denotes nodes present in an edge.

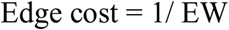

The main focus of the study was to identify the key players involved in regulating the variations in different conditions. We carried out shortest path analysis on the condition-specific networks and selected the paths that are most perturbed in these conditions. We implemented shortest path algorithm to obtain the results.

### Shortest path analysis

The edge cost values were used as an input for calculating all vs. all shortest paths in each condition using Zen (http://www.networkdynamics.org/static/zen/html/api/algorithms/shortest_path.html). More than 9000000 paths were obtained for each condition. In order to analyze the more significant paths, we ordered the paths on the basis of their path scores. Path score is the summation of the edge cost that constitutes a path. Based on the formula considered for calculating edge cost, lower path score indicates that the nodes in the path have higher expression. So, instead of analyzing 9000000 paths, we considered subnetworks, which comprise of top 1% of the network. These networks were visualized using Cytoscape 3 (48). Our response networks competently explain the perturbations in the system upon exposure to different situations such as AG treatment and/or disruption of whiB4. The networks were further co-related to graph-theory based methods and differentially regulated paths were recognized in each condition to construct sub-network for each condition (Supplementary Table S8).

### qRT-PCR analysis

Total RNA was isolated as described previously (6) and cDNA was synthesized (after DNase treatment) from 500 ng isolated RNA. Random oligonucleotide primers were used with iScript^TM^ Select cDNA Synthesis Kit (Bio-Rad) for cDNA synthesis. Gene specific primers (Supplementary Table S9) were selected for RT-PCR (CFX96 RT-PCR system, Bio-Rad) and iQ^TM^ SYBR Green Supermix (Bio-Rad) was used for gene expression analysis. In order to obtain meticulous expression levels, PCR expression was normalized and CFX Manager^TM^ software (Bio-Rad) was utilized for data analysis. Gene expression was normalized to *Mtb* 16S rRNA expression.

#### Electrophoretic Mobility Shift Assays (EMSA)

The histidine-tagged WhiB4 purification and generation of reduced or oxidized apo-WhiB4 was done as described previously (6). For EMSA assays, the promoter fragments of *whiB4, blaC,* and *blaR* (~150 bp upstream of translational start codon) were PCR amplified from the *Mtb* genome and the 5’ end was labelled using [γ-^32^P]-ATP labelled oligonucleotides by using T4 polynucleotide kinase (MBI Fermentas) as per the manufacturer’s instructions. Binding reactions were performed in 1X TBE buffer (89 mM Tris, 89 mM boric acid and 1 mM EDTA; pH 8.4) for 30 min and 5% polyacrylamide gel was used to resolve protein-DNA complexes. Gels were exposed to auto radiographic film and visualized via phosphoimaging (GE).

#### *In vitro* transcription assays

50 nM of DNA fragment containing *blaC* promoter and apo-WhiB4 (oxidized) or (reduced) were incubated in transcription buffer; 50 mM Tris HCl, (pH 8.0), 10 mM magnesium acetate, 100 μM EDTA, 100 μM DTT, 50 mM KCl, 50 μg/ml BSA, and 5% glycerol) for 30 minute at room temperature. Single round transcription assay was initiated by the addition of *Msm* RNAP-σ^A^ holo enzyme (100 nM), 100 µM NTPs and 1µCi α-^32^P-UTP and incubated at 37°C for 20 min. Reactions were terminated with 2X stop dye (95% formamide, 0.025 % (w/v) bromophenol blue, 0.025% (w/v) xylene cyanol, 5 mM EDTA and 0.025 % SDS and 8 M urea) and heated at 95°C for 5 minute followed by snap chilling in ice for 2 minutes. The transcripts were resolved by loading samples on to 6% urea-PAGE.

#### Intramycobacterial E_MSH_ measurement in vitro and during infection

Mycobacterial strains expressing Mrx1-roGFP2 were grown in 7H9 medium till an OD_600_ 0.4 and exposed to various concentration of AG. For measurements, cell were treated with 10 mM N-Ethylmaleimide (NEM) for 5 min at room temperature (RT) followed by fixation with 4% paraformaldehyde (PFA) for 15 min at RT. After washing thrice with 1X PBS, bacilli were analyzed using BD FACS Verse Flow cytometer (BD Biosciences). The biosensor response was measured by analyzing the ratio at a fixed emission (510/10 nm) after excitation at 405 and 488 nm as described (4). Data was analyzed using the FACSuite software. For measuring intramycobacterial *E_MSH_* during infection, PMA-differentiated THP-1 cells were infected with *Mtb* strains expressing Mrx1-roGFP2 (moi: 10). Infected macrophages were treated with NEM/PFA, washed with 1X PBS, and analyzed by flow cytometry as described previously (39).

#### Survival assay upon AG treatment in vitro and during infection

*Mtb* strains were grown aerobically till OD_600_ 0.4, followed by treatment with various concentrations of AG. At defined time-points post-exposure, cells were serially diluted and plated on OADC-7H11 agar medium for enumerating CFUs. To determine the effect of AG during infection, ~20000 THP-1 cells (PMA differentiated) were infected with wt *Mtb* in a 96-well plate (moi:10) as described earlier (39). Various concentrations of AG were added immediately after infection and at various time points macrophages were lysed using 0.06 % SDS-7H9 medium and released bacteria were serially diluted and plated on OADC-7H11 agar medium for CFU determination.

#### Vancomycin-BODIPY staining

The pattern of nascent peptidoglycan synthesis was observed by fluorescent staining as described (53). *Mtb* strains were grown to exponential phase (OD_600_ 0.6) in 7H9 medium. 1 ml of culture was incubated with 1 µg/ml Vancomycin-BODIPY (BODIPY^®^ FL Vancomycin, Thermo Fisher Scientific) for 16 h under standard growth conditions. Cells were pelleted to remove excess stain and fixed with PFA. After washing with 1X PBS, culture aliquots (20 µl) were spread on slides and allowed to air dry. The bacterial cells were visualized for BODIPY^®^ FL Vancomycin fluorescence (excitation at 560 nm and emission at 590 nm) in a Leica TCS Sp5 confocal microscope under a 63X oil immersion objective. Staining pattern of more than 150 cells was observed for each strain and cell length was measured as well using Image J software.

#### Aerosol infection of mice

For acute model of infection, BALB/c mice were infected via aerosol with 10,000 bacilli per mouse with the *Mtb H37Rv*, *Mtb∆whiB4, whiB4-OE,* MYC 431 and MYC 431-*whiB4-OE* strains as described (52). For assured over-expression of WhiB4, doxycycline was supplied, dissolved in drinking water; 1 mg/ml in 5% sucrose solution. Amox dose was maintained as 200 mg/kg of body weight while Clav dose was maintained as 50 mg/kg of body weight and drugs were administered orally twice a day. At specific time points, mice were sacrificed and their lungs were removed and processed for investigation of bacillary load. CFUs were determined by plating appropriate serial dilutions on 7H11(supplemented with OADC) plates. Colonies were observed and counted after 3-4 weeks of incubation at 37°C.

#### Statistical Analysis

Statistical analyses were performed using GraphPad Prism software. The statistical significance of the differences between experimental groups was determined by two-tailed, unpaired Student’s t test. Differences with a *p value* of ≤0.05 were considered significant.

#### Ethics statement

This study was carried out in strict accordance with the guidelines provided by the Committee for the Purpose of Control and Supervision on Experiments on Animals (CPCSEA), Government of India. The protocol was approved by the Committee on the Ethics of Animal Experiments of the International Centre for Genetic Engineering and Biotechnology (ICGEB), New Delhi, India (Approval number: ICGEB/AH/2011/2/IMM-26). All efforts were made to minimize the suffering.

## Acknowledgement

We are thankful to the University of Delhi South Campus MicroArray Centre (UDSCMAC), New Delhi for conducting microarray experiment. We are grateful to Dr. Harinath Chakrapani (IISER, Pune) and Dr. Santosh Podder (IISc, Bangalore) for excellent technical help with DHE assay and confocal microscopy, respectively. We thank Dr. William R. Jacobs, Jr. (Albert Einstein College of Medicine) for the *MtbΔmshA* and *mshA* complemented strains, David R. Sherman (Seattle Biomed, USA) for TetRO based *E.coli-mycobacterial* shuttle vector, *pEXCF-whiB4*, Dr. Y Av-Gay (University of British Columbia, Vancouver, British Columbia, Canada) for *MsmΔmshA* mutant. The *Mtb* work was supported by the Wellcome-DBT India Alliance grant, WT-DBT/500034-Z-09-Z (A.S.), and in part by Department of Biotechnology (DBT) Grant BT/PR5020/MED/29/1454/2012 (A.S.) and DBT-IISc program. A.S. is a Wellcome DBT India Alliance Intermediate Fellow. We acknowledge DBT-IISc-supported BSL3 facility for carrying out experiments on *Mtb* strains.

## References

1 Al-Attar S, Yu Y, Pinkse M, Hoeser J, Friedrich T, Bald D, de Vries S. Cytochrome bd Displays Significant Quinol Peroxidase Activity. Sci Rep 6: 27631, 2016.

2. Aly SA, Boothe DM, Suh SJ. A novel alanine to serine substitution mutation in SoxS induces overexpression of efflux pumps and contributes to multidrug resistance in clinical Escherichia coli isolates. J Antimicrob Chemother 70: 2228-33, 2015.

3. Baek SH, Li AH, Sassetti CM. Metabolic regulation of mycobacterial growth and antibiotic sensitivity. PLoS Biol 9: e1001065, 2011.

4. Bhaskar A, Chawla M, Mehta M, Parikh P, Chandra P, Bhave D, Kumar D, Carroll KS, Singh A. Reengineering redox sensitive GFP to measure mycothiol redox potential of Mycobacterium tuberculosis during infection. PLoS Pathog 10: e1003902, 2014.

5. Chambers HF, Kocagoz T, Sipit T, Turner J, Hopewell PC. Activity of amoxicillin/clavulanate in patients with tuberculosis. Clin Infect Dis 26: 874-7, 1998.

6. Chawla M, Parikh P, Saxena A, Munshi M, Mehta M, Mai D, Srivastava AK, Narasimhulu KV, Redding KE, Vashi N, Kumar D, Steyn AJ, Singh A. Mycobacterium tuberculosis WhiB4 regulates oxidative stress response to modulate survival and dissemination in vivo. Mol Microbiol 85: 1148-65, 2012.

7. Cruz-Ramos H, Cook GM, Wu G, Cleeter MW, Poole RK. Membrane topology and mutational analysis of Escherichia coli CydDC, an ABC-type cysteine exporter required for cytochrome assembly. Microbiology 150: 3415-27, 2004.

8. Cynamon MH, Palmer GS. In vitro activity of amoxicillin in combination with clavulanic acid against Mycobacterium tuberculosis. Antimicrob Agents Chemother 24: 429-31, 1983.

9. Deretic V, Philipp W, Dhandayuthapani S, Mudd MH, Curcic R, Garbe T, Heym B, Via LE, Cole ST. Mycobacterium tuberculosis is a natural mutant with an inactivated oxidative-stress regulatory gene: implications for sensitivity to isoniazid. Mol Microbiol 17: 889-900, 1995.

10. Dinesh N, Sharma S, Balganesh M. Involvement of efflux pumps in the resistance to peptidoglycan synthesis inhibitors in Mycobacterium tuberculosis. Antimicrob Agents Chemother 57: 1941-3, 2013.

11. Dwyer DJ, Belenky PA, Yang JH, MacDonald IC, Martell JD, Takahashi N, Chan CT, Lobritz MA, Braff D, Schwarz EG, Ye JD, Pati M, Vercruysse M, Ralifo PS, Allison KR, Khalil AS, Ting AY, Walker GC, Collins JJ. Antibiotics induce redox-related physiological alterations as part of their lethality. Proc Natl Acad Sci U S A 111: E2100-9, 2014.

12. Fay A, Glickman MS. An essential nonredundant role for mycobacterial DnaK in native protein folding. PLoS Genet 10: e1004516, 2014.

13. Fishbein S, van Wyk N, Warren RM, Sampson SL. Phylogeny to function: PE/PPE protein evolution and impact on Mycobacterium tuberculosis pathogenicity. Mol Microbiol 96: 901-16, 2015.

14. Flores AR, Parsons LM, Pavelka MS, Jr. Characterization of novel Mycobacterium tuberculosis and Mycobacterium smegmatis mutants hypersusceptible to beta-lactam antibiotics. J Bacteriol 187: 1892-900, 2005.

15. Flores AR, Parsons LM, Pavelka MS, Jr. Genetic analysis of the beta-lactamases of Mycobacterium tuberculosis and Mycobacterium smegmatis and susceptibility to beta-lactam antibiotics. Microbiology 151: 521-32, 2005.

16. Gold B, Pingle M, Brickner SJ, Shah N, Roberts J, Rundell M, Bracken WC, Warrier T, Somersan S, Venugopal A, Darby C, Jiang X, Warren JD, Fernandez J, Ouerfelli O, Nuermberger EL, Cunningham-Bussel A, Rath P, Chidawanyika T, Deng H, Realubit R, Glickman JF, Nathan CF. Nonsteroidal anti-inflammatory drug sensitizes Mycobacterium tuberculosis to endogenous and exogenous antimicrobials. Proc Natl Acad Sci U S A 109: 16004-11, 2012.

17. Gregory PD, Lewis RA, Curnock SP, Dyke KG. Studies of the repressor (BlaI) of beta-lactamase synthesis in Staphylococcus aureus. Mol Microbiol 24: 1025-37, 1997.

18. Gupta R, Lavollay M, Mainardi JL, Arthur M, Bishai WR, Lamichhane G. The Mycobacterium tuberculosis protein LdtMt2 is a nonclassical transpeptidase required for virulence and resistance to amoxicillin. Nat Med 16: 466-9, 2010.

19. Gusarov I, Shatalin K, Starodubtseva M, Nudler E. Endogenous nitric oxide protects bacteria against a wide spectrum of antibiotics. Science 325: 1380-4, 2009.

20. Harms A, Maisonneuve E, Gerdes K. Mechanisms of bacterial persistence during stress and antibiotic exposure. Science 354, 2016.

21. Hu Y, Coates AR. Acute and persistent Mycobacterium tuberculosis infections depend on the thiol peroxidase TpX. PLoS One 4: e5150, 2009.

22. Hugonnet JE, Tremblay LW, Boshoff HI, Barry CE, 3rd, Blanchard JS. Meropenem-clavulanate is effective against extensively drug-resistant Mycobacterium tuberculosis. Science 323: 1215-8, 2009.

23. Imlay JA. Pathways of oxidative damage. Annu Rev Microbiol 57: 395-418, 2003.

24. Imlay JA. The molecular mechanisms and physiological consequences of oxidative stress: lessons from a model bacterium. Nat Rev Microbiol 11: 443-54, 2013.

25. Kalyanaraman B, Dranka BP, Hardy M, Michalski R, Zielonka J. HPLC-based monitoring of products formed from hydroethidine-based fluorogenic probes--the ultimate approach for intra- and extracellular superoxide detection. Biochim Biophys Acta 1840: 739-44, 2014.

26. Kohanski MA, Dwyer DJ, Hayete B, Lawrence CA, Collins JJ. A common mechanism of cellular death induced by bactericidal antibiotics. Cell 130: 797-810, 2007.

27. Lamprecht DA, Finin PM, Rahman MA, Cumming BM, Russell SL, Jonnala SR, Adamson JH, Steyn AJ. Turning the respiratory flexibility of Mycobacterium tuberculosis against itself. Nat Commun 7: 12393, 2016.

28. Lobritz MA, Belenky P, Porter CB, Gutierrez A, Yang JH, Schwarz EG, Dwyer DJ, Khalil AS, Collins JJ. Antibiotic efficacy is linked to bacterial cellular respiration. Proc Natl Acad Sci U S A 112: 8173-80, 2015.

29. Lu P, Heineke MH, Koul A, Andries K, Cook GM, Lill H, van Spanning R, Bald D. The cytochrome bd-type quinol oxidase is important for survival of Mycobacterium smegmatis under peroxide and antibiotic-induced stress. Sci Rep 5: 10333, 2015.

30. Lun S, Miranda D, Kubler A, Guo H, Maiga MC, Winglee K, Pelly S, Bishai WR. Synthetic lethality reveals mechanisms of Mycobacterium tuberculosis resistance to beta-lactams. MBio 5: e01767-14, 2014.

31. Maksymiuk C, Balakrishnan A, Bryk R, Rhee KY, Nathan CF. E1 of alpha-ketoglutarate dehydrogenase defends Mycobacterium tuberculosis against glutamate anaplerosis and nitroxidative stress. Proc Natl Acad Sci U S A 112: E5834-43, 2015.

32. Master SS, Springer B, Sander P, Boettger EC, Deretic V, Timmins GS. Oxidative stress response genes in Mycobacterium tuberculosis: role of ahpC in resistance to peroxynitrite and stage-specific survival in macrophages. Microbiology 148: 3139-44, 2002.

33. Mehta M, Rajmani RS, Singh A. Mycobacterium tuberculosis WhiB3 Responds to Vacuolar pH-induced Changes in Mycothiol Redox Potential to Modulate Phagosomal Maturation and Virulence. J Biol Chem 291: 2888-903, 2016.

34. Miller C, Thomsen LE, Gaggero C, Mosseri R, Ingmer H, Cohen SN. SOS response induction by beta-lactams and bacterial defense against antibiotic lethality. Science 305: 1629-31, 2004.

35. Mir M, Asong J, Li X, Cardot J, Boons GJ, Husson RN. The extracytoplasmic domain of the Mycobacterium tuberculosis Ser/Thr kinase PknB binds specific muropeptides and is required for PknB localization. PLoS Pathog 7: e1002182, 2011.

36. Morris RP, Nguyen L, Gatfield J, Visconti K, Nguyen K, Schnappinger D, Ehrt S, Liu Y, Heifets L, Pieters J, Schoolnik G, Thompson CJ. Ancestral antibiotic resistance in Mycobacterium tuberculosis. Proc Natl Acad Sci U S A 102: 12200-5, 2005.

37. Nandakumar M, Nathan C, Rhee KY. Isocitrate lyase mediates broad antibiotic tolerance in Mycobacterium tuberculosis. Nat Commun 5: 4306, 2014.

38. Nguyen D, Joshi-Datar A, Lepine F, Bauerle E, Olakanmi O, Beer K, McKay G, Siehnel R, Schafhauser J, Wang Y, Britigan BE, Singh PK. Active starvation responses mediate antibiotic tolerance in biofilms and nutrient-limited bacteria. Science 334: 982-6, 2011.

39. Padiadpu J, Baloni P, Anand K, Munshi M, Thakur C, Mohan A, Singh A, Chandra N. Identifying and Tackling Emergent Vulnerability in Drug-Resistant Mycobacteria. ACS Infect Dis 2: 592-607, 2016.

40. Pathania R, Navani NK, Gardner AM, Gardner PR, Dikshit KL. Nitric oxide scavenging and detoxification by the Mycobacterium tuberculosis haemoglobin, HbN in Escherichia coli. Mol Microbiol 45: 1303-14, 2002.

41. Rao SP, Alonso S, Rand L, Dick T, Pethe K. The protonmotive force is required for maintaining ATP homeostasis and viability of hypoxic, nonreplicating Mycobacterium tuberculosis. Proc Natl Acad Sci U S A 105: 11945-50, 2008.

42. Rice KC, Firek BA, Nelson JB, Yang SJ, Patton TG, Bayles KW. The Staphylococcus aureus cidAB operon: evaluation of its role in regulation of murein hydrolase activity and penicillin tolerance. J Bacteriol 185: 2635-43, 2003.

43. Rullas J, Dhar N, McKinney JD, Garcia-Perez A, Lelievre J, Diacon AH, Hugonnet JE, Arthur M, Angulo-Barturen I, Barros-Aguirre D, Ballell L. Combinations of beta-Lactam Antibiotics Currently in Clinical Trials Are Efficacious in a DHP-I-Deficient Mouse Model of Tuberculosis Infection. Antimicrob Agents Chemother 59: 4997-9, 2015.

44. Saini V, Cumming BM, Guidry L, Lamprecht DA, Adamson JH, Reddy VP, Chinta KC, Mazorodze JH, Glasgow JN, Richard-Greenblatt M, Gomez-Velasco A, Bach H, Av-Gay Y, Eoh H, Rhee K, Steyn AJ. Ergothioneine Maintains Redox and Bioenergetic Homeostasis Essential for Drug Susceptibility and Virulence of Mycobacterium tuberculosis. Cell Rep 14: 572-85, 2016.

45. Sala C, Haouz A, Saul FA, Miras I, Rosenkrands I, Alzari PM, Cole ST. Genome-wide regulon and crystal structure of BlaI (Rv1846c) from Mycobacterium tuberculosis. Mol Microbiol 71: 1102-16, 2009.

46. Sambarey A, Prashanthi K, Chandra N. Mining large-scale response networks reveals 'topmost activities' in Mycobacterium tuberculosis infection. Sci Rep 3: 2302, 2013.

47. Sambaturu N, Mishra M, Chandra N. EpiTracer - an algorithm for identifying epicenters in condition-specific biological networks. BMC Genomics 17 Suppl 4: 543, 2016.

48. Shannon P, Markiel A, Ozier O, Baliga NS, Wang JT, Ramage D, Amin N, Schwikowski B, Ideker T. Cytoscape: a software environment for integrated models of biomolecular interaction networks. Genome Res 13: 2498-504, 2003.

49. Shatalin K, Shatalina E, Mironov A, Nudler E. H2S: a universal defense against antibiotics in bacteria. Science 334: 986-90, 2011.

50. Singh A, Crossman DK, Mai D, Guidry L, Voskuil MI, Renfrow MB, Steyn AJ. Mycobacterium tuberculosis WhiB3 maintains redox homeostasis by regulating virulence lipid anabolism to modulate macrophage response. PLoS Pathog 5: e1000545, 2009.

51. Soetaert K, Rens C, Wang XM, De Bruyn J, Laneelle MA, Laval F, Lemassu A, Daffe M, Bifani P, Fontaine V, Lefevre P. Increased Vancomycin Susceptibility in Mycobacteria: a New Approach To Identify Synergistic Activity against Multidrug-Resistant Mycobacteria. Antimicrob Agents Chemother 59: 5057-60, 2015.

52. Solapure S, Dinesh N, Shandil R, Ramachandran V, Sharma S, Bhattacharjee D, Ganguly S, Reddy J, Ahuja V, Panduga V, Parab M, Vishwas KG, Kumar N, Balganesh M, Balasubramanian V. In vitro and in vivo efficacy of beta-lactams against replicating and slowly growing/nonreplicating Mycobacterium tuberculosis. Antimicrob Agents Chemother 57: 2506-10, 2013.

53. Thanky NR, Young DB, Robertson BD. Unusual features of the cell cycle in mycobacteria: polar-restricted growth and the snapping-model of cell division. Tuberculosis (Edinb) 87: 231-6, 2007.

54. Tomasz A. The role of autolysins in cell death. Ann N Y Acad Sci 235: 439-47, 1974.

55. Tyagi P, Dharmaraja AT, Bhaskar A, Chakrapani H, Singh A. Mycobacterium tuberculosis has diminished capacity to counteract redox stress induced by elevated levels of endogenous superoxide. Free Radic Biol Med 84: 344-54, 2015.

56. Ung KS, Av-Gay Y. Mycothiol-dependent mycobacterial response to oxidative stress. FEBS Lett 580: 2712-6, 2006.

57. Vaubourgeix J, Lin G, Dhar N, Chenouard N, Jiang X, Botella H, Lupoli T, Mariani O, Yang G, Ouerfelli O, Unser M, Schnappinger D, McKinney J, Nathan C. Stressed mycobacteria use the chaperone ClpB to sequester irreversibly oxidized proteins asymmetrically within and between cells. Cell Host Microbe 17: 178-90, 2015.

58. Venugopal A, Bryk R, Shi S, Rhee K, Rath P, Schnappinger D, Ehrt S, Nathan C. Virulence of Mycobacterium tuberculosis depends on lipoamide dehydrogenase, a member of three multienzyme complexes. Cell Host Microbe 9: 21-31, 2011.

59. Vilcheze C, Hartman T, Weinrick B, Jacobs WR, Jr. Mycobacterium tuberculosis is extraordinarily sensitive to killing by a vitamin C-induced Fenton reaction. Nat Commun 4: 1881, 2013.

60. Wang X, Zhao X. Contribution of oxidative damage to antimicrobial lethality. Antimicrob Agents Chemother 53: 1395-402, 2009.

